# Salicylic acid-triggered apoplastic proteolysis releases cryptic phytocytokines with distinct immunogenic functions

**DOI:** 10.64898/2026.09.01.748544

**Authors:** Philipp Katzy, Hermanyanto Laia, Zoé Zick, Johana Misas Villamil, Gunther Doehlemann

**Affiliations:** Institute for Plant Sciences and Cluster of Excellence on Plant Sciences (CEPLAS), University of Cologne, Cologne, German

**Keywords:** Phytocytokines, apoplast, plant immune signalling, plant proteases, cryptic peptides, *Zea mays*

## Abstract

Plants rely on an innate immune system to defend against pathogens through various molecular responses. In addition to classical damage-and pathogen-associated molecular patterns (DAMPs and PAMPs), plants produce endogenous signaling peptides termed phytocytokines that amplify and regulate immune responses following stress. Although most characterized phytocytokines originate from dedicated precursor proteins, the contribution of multifunctional proteins to phytocytokine generation remains poorly understood. Here, we show that salicylic acid (SA) rapidly remodels the maize apoplastic peptidome through an early, transient proteolytic program driven by apoplastic serine hydrolases. Time course peptidomics identified fourteen candidate phytocytokines, including two cryptic peptides, PC13 and PC14, released from the stress-associated zincfinger protein ZmSAP7 and the migration inhibitory factor-like protein ZmMDL1, respectively. Both peptides activated immune-associated gene expression but triggered distinct transcriptional responses and exerted opposing effects on *Ustilago maydis* infection, with PC13 enhancing resistance and PC14 promoting susceptibility. Biochemical analysis demonstrated that PMSF-sensitive apoplastic serine proteases directly process ZmMDL1 to release PC14. Together, our findings uncover a SA-responsive proteolytic pathway that generates functionally distinct phytocytokines from multifunctional proteins, expanding the repertoire of immune signaling peptides and revealing an additional layer of regulation in plant defense.

## Introduction

Plants have evolved a highly complex immune system that enables them to defend against a wide range of pathogens, including viruses, bacteria, fungi, oomycetes, herbivores, and parasitic plants (Ngou et al., 2022). The first barrier encountered by invading pathogens is the plant cell wall, which may be breached through wounding, enzymatic degradation, or by entering through stomata. Once pathogens overcome this barrier, they encounter the plasma membrane, where pattern recognition receptors (PRRs) detect conserved molecular signatures of microbes, termed microbe-or pathogen-associated molecular patterns (MAMPs/PAMPs) (Peng et al., 2018). PRRs also recognize endogenous signals derived from plant tissue damage (DAMPs), forming the basis of pattern-triggered immunity (Boller & Felix, 2009; Ma et al., 2024; Ngou et al., 2022). Perception of these signals induces a broad spectrum of immune responses, including Ca²⁺ influx, reactive oxygen species (ROS) production, mitogen-activated protein kinase (MAPK) activation, and biosynthesis of salicylic acid (SA) and jasmonic acid (JA) (Lecourieux et al., 2002; Meng & Zhang, 2013; Rodríguez et al., 2006).

In recent years, PRRs have also been shown to perceive plant-derived peptides, termed phytocytokines, which act as endogenous regulators of immunity and development (Hou et al., 2021). Initially classified as DAMPs, phytocytokines differ in that their release is tightly regulated, requiring proteolytic cleavage of canonical peptide precursor proteins whereas classical DAMPs such as extracellular ATP or oligogalacturonides are passively released upon damage (Choi & Klessig, 2016; Gust et al., 2017). Phytocytokines can originate from two major classes of precursors: non-functional precursor proteins, which give rise to peptides such as post-translationally modified peptides, cysteine-rich peptides, or unmodified peptides lacking cysteine residues and modifications; and functional precursor proteins, which possess an independent biological activity beyond peptide generation (S. Wang et al., 2026). Although most phytocytokines are encoded as dedicated precursor proteins, a growing number of examples demonstrate that functional proteins can also harbor cryptic peptide signals that are released upon proteolysis and contribute to plant immune responses. These include CAPE1, derived from the pathogenesis-related protein PR1b, GmSUBPEP, embedded within a soybean subtilase, and inceptins, which are generated from the chloroplastic ATP synthase γ-subunit. (Castaldi et al., 2026; Pearce et al., 2010a; Schmelz et al., 2006).

Phytocytokines are also classified according to whether their canonical precursor protein contain an N-terminal signal peptide (SP) for secretion or not. SP-containing precursors are secreted via the canonical ER–Golgi pathway and undergo additional processing to yield active peptides, as shown for HypSys, PIP1/2, PSK, PSY1, IDA, and RALFs (Amano et al., 2007; Butenko, 2003; Hou et al., 2014; Narváez-Vásquez et al., 2005; Pearce et al., 2001, 2010b). In contrast, precursors lacking a signal peptide may follow unconventional secretion pathways, in which non-secreted precursor proteins undergo a two-step maturation process involving both intracellular and extracellular proteolytic cleavage (Koenig et al., 2026; S. Wang et al., 2026). Otherwise, peptides originated from precursor proteins lacking a signal peptide are believed to be released upon cellular damage, with maturation occurring in the cytosol or apoplast. For example, the peptide PEP1 is generated upon damage from the signal peptide-lacking precursor PROPEP1 by metacaspases MC4–MC9 in *A. thaliana* (Hander et al., 2019; Shen et al., 2019). Proteases are therefore key regulators of phytocytokine maturation and activity. The wheat papain-like cysteine protease (PLCPs) *Ta*RD21A releases the small signaling peptide Wip1, which confers wheat resistance to wheat yellow mosaic virus (Liu et al., 2024). In maize, the Zip1 phytocytokine was identified in the maize apoplast 24 hours after salicylic acid (SA) treatment. Zip1 functions as amplifier of SA-dependent immune responses by activating apoplastic PLCPs (Ziemann et al., 2018). The maize PLCP immune proteases CP1 and CP2 orchestrate processing of PROZIP1 to confer release, as well as clearance of the mature Zip1 in the apoplast. However, before apoplastic maturation, PROZIP1 undergoes a first intracellular processing performed by the metacaspase *Zm*MC9 (Koenig et al., 2026). Subtilases (SBTs) also contribute to phytocytokine processing and broader immune regulation. In *A. thaliana*, for instance, SBT4.12, SBT4.13, and SBT5.2 redundantly cleave proIDA to release IDA which regulates cell separation and modulates plant immunity (Lalun et al., 2024; Schardon et al., 2016). In tomato, *Sl*Phyt1 and *Sl*Phyt2 release PSK and Systemin from their propeptides (Beloshistov et al., 2018; Reichardt et al., 2020).

In this study, we conducted time-resolved profiling of proteolytic activities in maize leaves upon SA-triggered immune induction, which revealed a transient early activation of serine hydrolase activity. From apoplastic fluids of SA-induced leaves, we identified native peptides, including two cryptic phytocytokines, PC13 and PC14, which are derived from functional proteins and mediate contrasting immune responses in maize leaves. Together, this work reveals temporal dynamics of SA-signaling and links transient subtilase activation with the release of previously unknown immunogenic signaling peptides released from functional precursor proteins, thereby expanding our understanding of peptide-mediated immune regulation.

## Results

### Dynamic release of immunogenic apoplastic peptides upon SA treatment

The maize phytocytokine Zip1 was identified in the maize apoplast 24 hours after SA treatment (Ziemann et al., 2018). Since immune signaling processes are highly dynamic and the 24 h timepoint likely reflects a late stage of SA signaling, we aimed to investigate the temporal dynamics of peptide release during the early response to SA. We performed an experiment in which maize leaves were infiltrated with either SA or a mock solution (see methods for details) and apoplastic fluid (AF) samples were collected at different time points. AF was filtered using a 10 kDa cutoff ultra-centrifugal device to collect the low molecular fraction, containing peptides, at 3, 6, 12, and 24 h after treatment. Each apoplastic fluid fraction (APF) was subsequently re-infiltrated into the second leave of naive 8-day-old maize seedlings. The transcriptional response of SA-responsive genes such as *PRm6b* and *PR10* was monitored 24 h after treatment (**Fig. 1A**).

**Figure 1:**
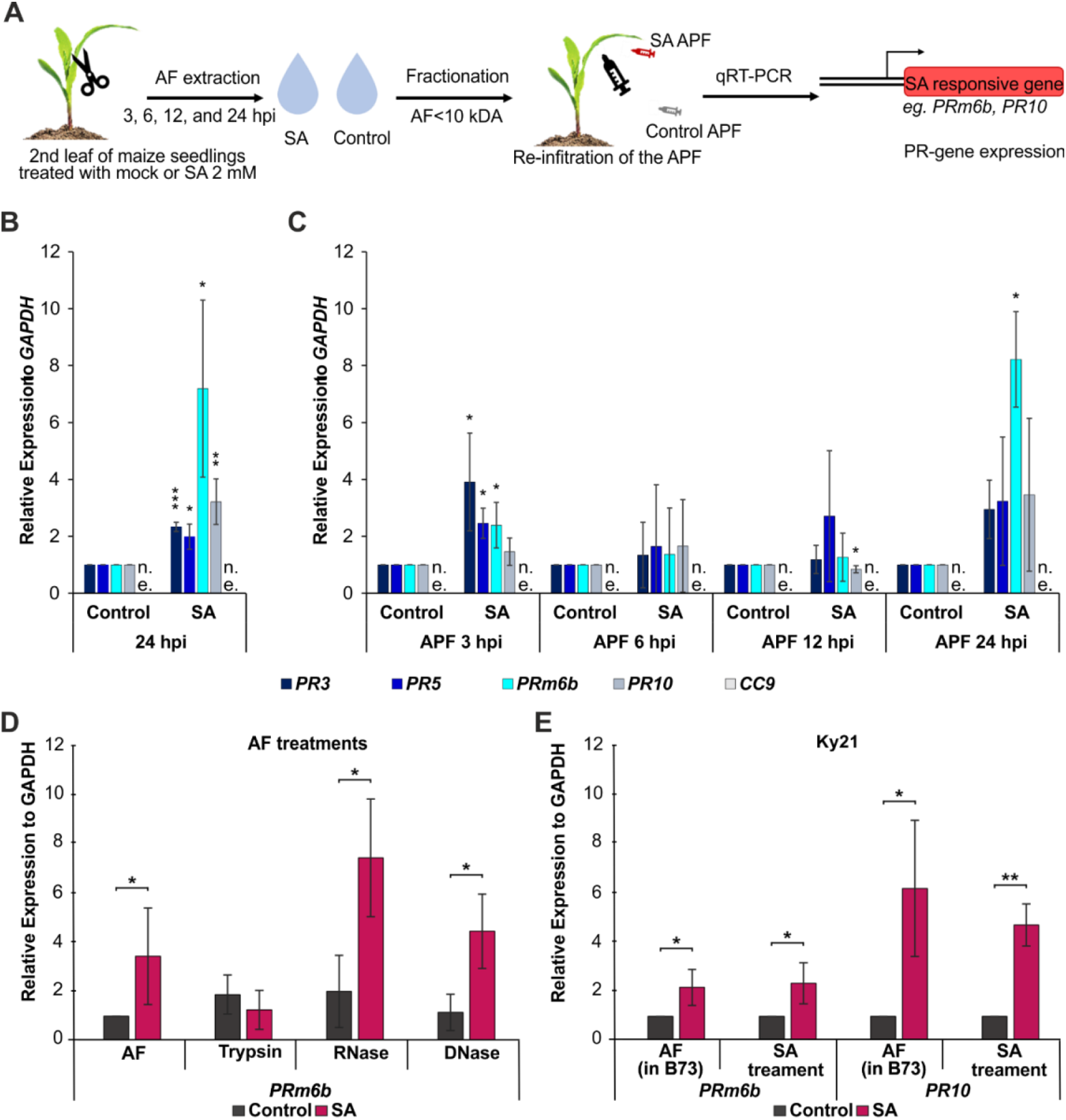
Peptides isolated from the SA-treated apoplastic fraction induce PR gene expression. (A) Schematic representation of the experimental procedure. Apoplastic fluid (AF) of mock and SA treated leaves was isolated at different time points. AF was filtered with a 10 kDa ultrafiltration device and the flow-through was reinfiltrated with a syringe into second leaves of 8-day old plants. Expression of *PR*-genes and *CC9* (JA marker) was determined 24 h after infiltration. (B-C) SA treatment induces PR gene expression in newly infiltrated leaves. Relative expression of PR3, PR5, PRm6b, PR10, and CC9 was quantified 24 h after SA or mock infiltration and following re-infiltration of peptide fractions isolated from the corresponding apoplastic fluids. *CC9* was not expressed (n.e.). (D) Protease, but not nuclease, treatment abolishes the PR-inducing activity of the SA-derived apoplastic peptide fraction. Peptide fractions were treated with trypsin, RNase, or DNase, re-filtered to remove the enzymes, and re-infiltrated into B73 maize leaves. PRm6b expression was quantified 24 h after infiltration. (E) SA-triggered peptides induce PR gene expression independently of ZIP1. Leaves of maize Ky21, which lacks proZIP1, were infiltrated with 2 mM SA or mock solution. Peptide fractions were isolated from the apoplastic fluid 3 h after treatment and re-infiltrated into B73 leaves. PRm6b and PR10 expression was quantified 24 h after infiltration. Statistically significant differences were determined based on students t-test (* =p<0.05, ** =p<0.01, *** =p<0.001).

Confirming previous reports, SA treatment triggered a clear induction of multiple PR marker genes after 24 h compared to mock treatments (**Fig. 1B**), validating the effectiveness of the experimental setup. Importantly, expression of *CC9*, a marker for jasmonic acid (JA) and wounding responses (van der Linde et al., 2012), was not induced by SA treatment, suggesting that wounding did not contribute to the observed *PR* gene induction (**Fig. 1B**). When, APFs from different timepoints were re-infiltrated into naive leaves, SA-induced APFs triggered a significant upregulation of *PR* gene expression compared to mock, suggesting that biologically active peptides are released into the apoplast after SA perception (**Fig. 1C**). APFs collected at 3 h from SA-treated plants significantly induced the expression of *PR3*, *PR5*, and *PRm6b*, while *PR10* was not significantly altered. In contrast, APFs from 6 h showed no induction of any of the tested PR marker genes (**Fig. 1C**). At later timepoints, a different pattern was observed: APFs from 12 h significantly induced *PR10* whereas APFs from 24 h significantly induced *PRm6b* (**Fig. 1C**). These results indicate that the release of signaling peptides into the apoplast is both dynamic and time-dependent, with distinct waves of activity occurring at early and later phases of the SA response.

To determine the biochemical nature of the active elicitors, the 3 h APF was treated with proteases and nucleases such as RNase, DNase, or trypsin, followed by removal of residual enzymes through filtration and then a re-infiltration of the low molecular weight fraction into naive maize seedlings. qPCR analysis of *PRm6b* revealed that trypsin treatment, but not RNase and DNase treatments, abolished the ability of the APF to induce *PR* gene expression (**Fig. 1D**). This suggests that proteinaceous elicitors, specifically peptides, are responsible for the observed induction of *PR* gene expression. Given that Zip1 is known to be released into the maize apoplast following SA treatment (Koenig et al., 2023, 2026; Ziemann et al., 2018), we next tested whether the observed activity could be explained solely by Zip1. To address this, we performed the same experiment using the maize inbred line Ky21, which lacks the *PROZIP1* gene and therefore cannot produce Zip1 (Depotter et al., 2022). APF collected from SA-treated Ky21 plants at 3 h was re-infiltrated into maize B73 seedlings and tested for *PR* gene induction. Remarkably, this APF still induced expression of *PR5* and *PRm6b*, suggesting that additional, previously unknown SA-responsive peptides besides Zip1 contribute to *PR* gene activation by the APF (**Fig. 1E**).

To identify candidate peptides responsible for the observed signaling activity, we performed native peptidomic profiling of APFs collected from SA-and mock-treated maize leaves at multiple time points. The obtained peptide profiles differed markedly between SA-and mock-treated samples (**Fig. S1A**). A core set of 34.8% of SA-enriched peptides was detected at 6, 9 and 12 h time points, while the fraction of detected peptides significantly enriched after SA treatment increased over time from 12.7% to 16.7% (**Fig. S1B**). Because *PR* gene induction was already detected 3 h after SA treatment, we focused our subsequent analyses on this time point. Across all samples, a total of 4876 peptides were detected (**Table S1, Fig. 2A**). We found half of the peptides (∼57%) shared between the control and the SA treatment, whereas 1048 peptides were exclusively found in the SA sample and 1042 only in the control sample (**Fig. 2A**). Based on the peptide intensities and the significance the vast majority (∼92%) exhibited no significant difference between SA and control treatments (**Fig. 2B, black dots**). Besides, 195 peptides were significantly enriched following SA treatment (**Fig. 2B, magenta dots**), including 181 exclusively detected in SA samples (**Table S1**). Conversely, 200 peptides were enriched in the control APF (**Fig. 2B, blue dots**), with 190 of these unique to the control sample (**Table S1**). Most peptides induced by SA at 3 h persisted throughout the time course, with only a small subset showing decreased abundance over time (**Fig. S2C**).

**Figure 2:**
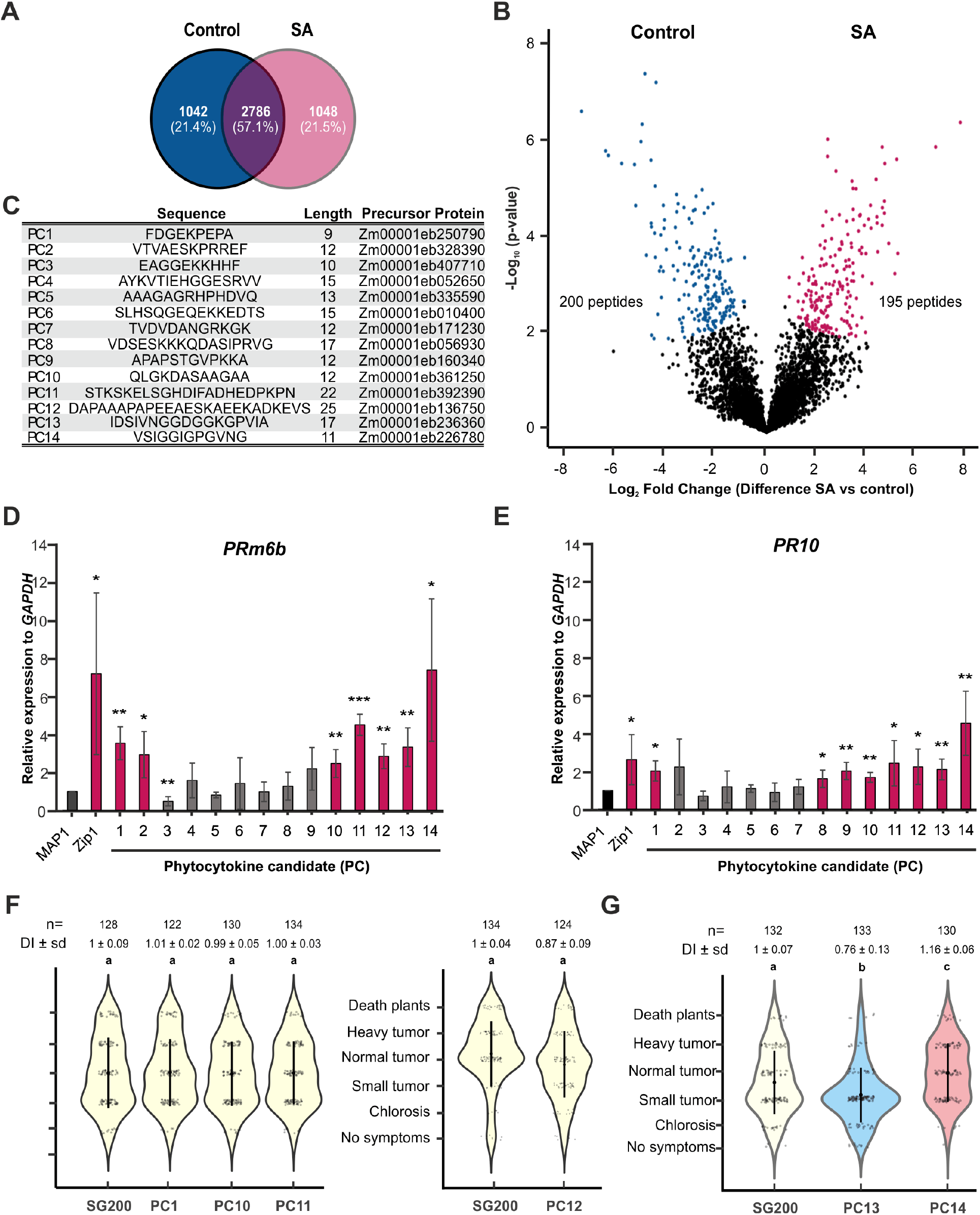
Two phytocytokine candidates induce *PR* gene expression and modulate *U. maydis* virulence. Second leaves from 8-day old maize plants were vacuum infiltrated with SA or DMSO (control) and AF was extracted at 3 h after treatment. The AF was filtered with a 10 kDa ultrafiltration device and the flow-through was used for peptidomic MS analysis. (A) Venn diagram of peptides identified 3 h after SA or mock treatment in at least three out of four replicates in either treatment. (B) Volcano plot of proteomic samples illustrating the distribution of the peptides within the treatments. Magenta and blue squares indicate significant differences (p-value <0.05, s0 = 0.1). (C) List of selected phytocytokine candidates (PC1-PC14) showing their peptide sequence, length and identified precursor protein. (D-E) Six phytocytokine candidates (PCs) consistently induce PR gene expression. Synthetic PCs (4 µM) were syringe-infiltrated into maize leaves. MAP1 and ZIP1 served as negative and positive controls, respectively. Leaf tissue was harvested 24 h after infiltration, and *PRm6b* (D) and *PR10* (E) transcript levels were quantified. Expression was normalized to GAPDH. PCs that significantly induced the respective PR gene are highlighted in purple, whereas those that did not are shown in grey. Significant differences were calculated based on students t-test (* =p<0.05, ** =p<0.01, *** =p<0.001). (F-G) *U. maydis*-secreted PC13 and PC14 differentially modulate fungal virulence. Seven-day-old maize seedlings were infected with *U. maydis* strain SG200 or transgenic SG200 strains expressing PC1, PC10, PC11, PC12, PC13, or PC14. Disease severity was quantified at 12 days post-infection (dpi). The disease index (DI) represents the mean symptom score relative to the wild-type SG200 strain, with symptoms scored from 0 (no symptoms) to 10 (dead plants). Statistical significance was determined by one-way ANOVA followed by Tukey’s HSD test, and plots were colored according to statistical groups. The number of analyzed plants is indicated by *n*. Data represent three independent biological replicates.

### Peptidomic profiling identifies SA-induced candidate phytocytokines

To prioritize candidate peptides for subsequent functional characterization, a series of filtering criteria was applied: i) peptides should be detected in at least three of four biological replicates of the SA treatment. ii) they need to be significantly more abundant in SA compared to control. iii) candidate peptides had to map uniquely to a single precursor protein. iv) peptides longer than 25 amino acids, as well as peptides mapping to universal motifs such as signal peptides were excluded. Finally, we prioritized candidates derived either from proteins with established roles in stress responses or from proteins lacking annotated domains, thereby representing both functional and putative dedicated precursor proteins. Following this pipeline, 14 phytocytokine candidates (PC1– PC14) were identified for further validation (**Fig. 2C**; **Table S3**). PC10, PC13 and PC14 were included because their precursor proteins have known or suspected roles in stress responses and could act as cryptic peptides. Besides, PC14 was detected at multiple timepoints (6, 9 and 12 h) only in SA-treated samples, suggesting sustained release (**Table S2**). Notably, the PC14 pro-protein was annotated as ZmMDL1 (Zm00001eb226780), a member of the macrophage migration inhibitory factor (MIF) family, which is well-characterized as a proinflammatory cytokine in mammals (Sumaiya et al., 2022) (**Table S3**).

To test whether the identified candidates function as bioactive signaling peptides, all 14 candidates were chemically synthesized and infiltrated into maize leaves. Induction of the defense marker genes *PRm6b* (**Fig. 2D**) and *PR10* (**Fig. 2E**) was quantified 24 h after infiltration. MAP1, an apoplastic peptide previously identified as accumulating in response to SA but lacking PR gene-inducing activity, served as a negative control, whereas Zip1 served as a positive control(Koenig et al., 2023; Ziemann et al., 2018). Six candidate peptides (PC1, PC10, PC11, PC12, PC13, and PC14) significantly induced the expression of both *PRm6b* and *PR10* (**Fig. 2D, E**). These findings suggest that several newly identified peptides may function as phytocytokines. To investigate the functional roles of candidate phytocytokines during a biotrophic interaction, we employed the “Trojan horse” approach, in which the maize pathogen *Ustilago maydis* is engineered to secrete candidate peptides into the plant apoplast during infection (van der Linde et al., 2018; Ziemann et al., 2018; Koenig et al., 2026). The six peptides PC1, PC10, PC11, PC12, PC13, and PC14 were expressed in *U. maydis*. Maize seedlings were inoculated with the recombinant strains, and disease symptoms were scored at 12 days post infection (dpi) relative to the progenitor strain SG200. Strains secreting PC1, PC10, PC11 and PC12 displayed no significant differences in virulence compared with SG200 (**Fig. 2F**). In contrast, secretion of PC13 and PC14 resulted in significant but opposing effects on disease development (**Fig. 2G**). PC13 significantly reduced *U. maydis* virulence, as indicated by smaller tumors and reduced chlorosis, consistent with a role in promoting plant resistance. Conversely, PC14 secretion resulted in a hypervirulent phenotype, with infected plants developing larger tumors compared with the SG200 control (**Fig. 2G**). Thus, although both PC13 and PC14 induced *PR* gene expression when applied as synthetic peptides, their effects diverged in the plant–pathogen interaction, revealing distinct functional outcomes of peptide-mediated immune modulation. The Trojan horse assay suggests that PC13 may function as a defense-promoting peptide, whereas PC14 may be exploited by *U. maydis* to enhance virulence, potentially through modulation of host signaling pathways that support biotrophic colonization.

### PC13 and PC14 trigger distinct and dynamic transcriptional programs

The opposing effects of PC13 and PC14 on *U. maydis* virulence prompted us to investigate their underlying host transcriptional responses. Transcriptome profiling was performed after treatment with synthetic PC13 and PC14 peptides at 2, 6, 12, and 24 h after infiltration (**Fig. 3, Table S4**). MAP1 served as a negative control, while Zip1 was included as a reference phytocytokine. Principal component analysis (PCA) revealed a temporal separation of transcriptomic responses, with samples collected at 2 h clustering distinctly from later time points along PC1 (**Fig. 3A**). Samples collected at 6, 12, and 24 h formed progressively separated clusters, reflecting dynamic changes in gene expression over time. Differential expression analysis revealed that all three peptides triggered the strongest transcriptional responses at 2 h after treatment (**Fig. 3B**). The number of up-regulated genes declined markedly by 6 h and reached a minimum at 12 h, followed by a second increase at 24 h, indicating a biphasic transcriptional response. Downregulated genes followed a similar temporal pattern, with the largest number detected at 2 h after treatment. PC14 caused the strongest repression, whereas PC13 and Zip1 resulted in fewer downregulated genes (**Fig. 3B, C**).

**Figure 3.**
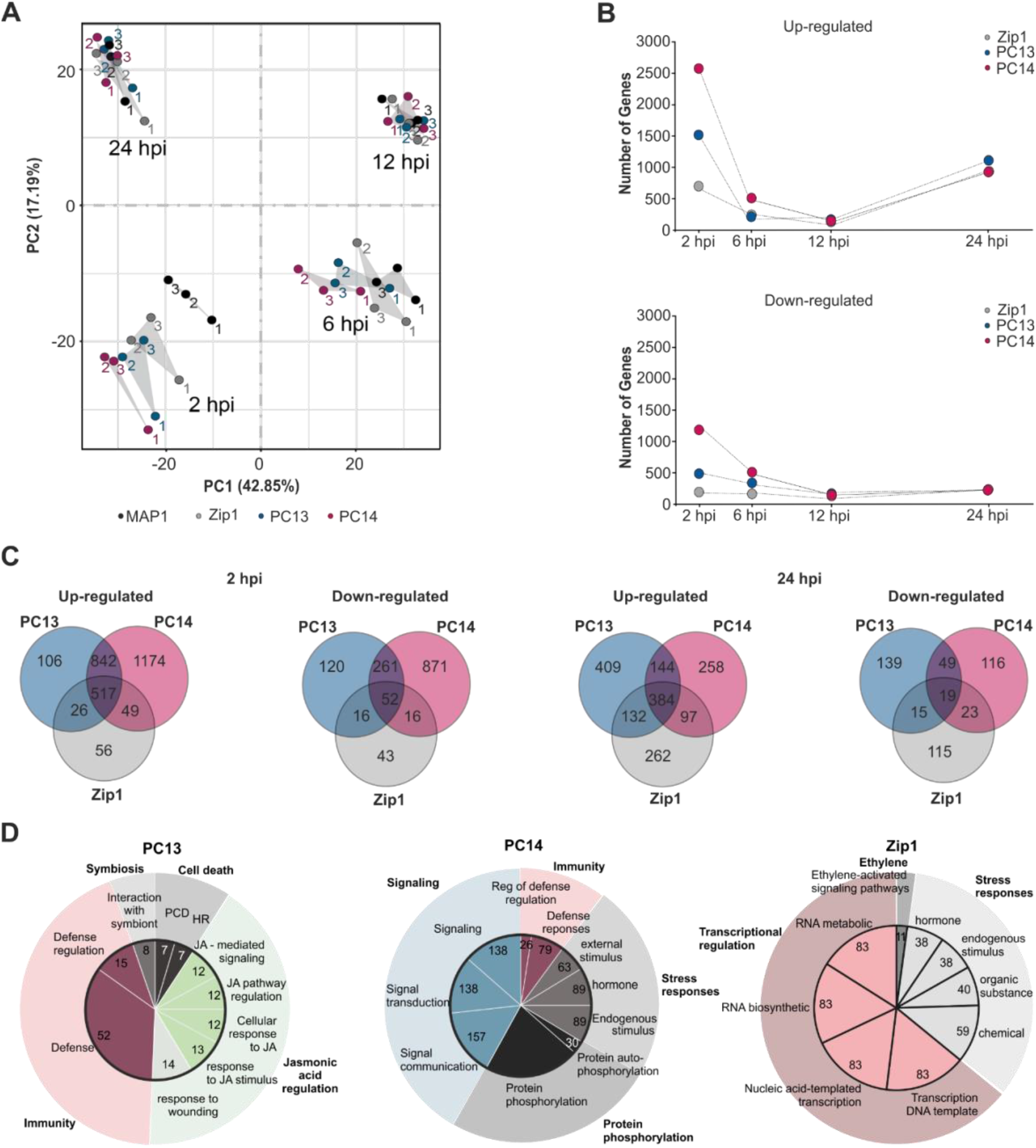
SA-induced phytocytokines trigger distinct early transcriptional immune responses in maize leaves. Seven-day-old maize seedlings were syringe-infiltrated with 4 µM synthetic Zip1, PC13, or PC14 peptides, with MAP1 serving as a negative control. Tissue surrounding the infiltration site was harvested for RNA sequencing at 2, 6, 12, and 24 h after treatment. (A) Principal component analysis (PCA) illustrates variation in gene expression among treatments and time points. (B) Number of differentially expressed genes (DEGs) significantly up-or downregulated following PC13, PC14, or Zip1 treatment. (C) Venn diagrams show treatment-specific and shared DEGs at 2 and 24 h after peptide treatment. Differential expression was determined using Student’s t-test (p < 0.05) with thresholds of log2 fold change > 1 or < −1 for up-and downregulated genes, respectively. (D) Gene Ontology (GO) enrichment analysis highlights biological processes associated with transcriptional responses to PC13, PC14, and Zip1 at 2 h after treatment.

Functional classification of genes up-regulated by PC13 at 2 h revealed enrichment of biological processes associated with plant defense and stress responses (**Fig. 3D, Table S5**). The largest categories included defense response, defense regulation, and response to wounding, indicating activation of immune-related pathways. Genes involved in jasmonic acid signaling and biosynthesis were also enriched. In addition, categories associated with symbiosis, programmed cell death, and hypersensitive response, were also represented, suggesting that PC13 induces a broad defense-related transcriptional response that may contribute to the reduced virulence phenotype observed in the Trojan horse assay (**Fig. 2G**). In contrast, genes up-regulated by PC14 were predominantly associated with signaling, protein phosphorylation processes and stress responses. Major enriched categories included protein phosphorylation, cell-to-cell communication, signal transduction, and signaling pathways. Defense-associated categories, including regulation of defense response, and responses to biotic stimulus, were also enriched although signaling-associated processes represented a larger proportion of PC14 transcriptional response (**Fig. 3D, Table S5**). Zip1 induced a broad transcriptional response characterized by enrichment of stress-and defense-associated categories, including ethylene-related signaling (**Table S5**). Together, these results indicate that the newly identified peptides PC13 and PC14, although both capable of inducing *PR* gene expression, engage distinct transcriptional programs. PC13 preferentially activates defense-and jasmonate-associated responses, whereas PC14 primarily induces signaling, modulation of stress responses and phosphorylation-related processes, reflecting functional diversification among SA-induced apoplastic peptides.

### PC13 and PC14 are cryptic phytocytokines derived from functional proteins

Unlike most known phytocytokines, both PC13 and PC14 originate from conserved maize proteins with established cellular functions. Thus, PC13 and PC14 represent cryptic phytocytokines, *i.e.* hidden signaling peptides embedded within functional proteins that provide an additional layer of immune regulation upon release. PC13 (IDSIVNGGDGGKGPVIA) is derived from the Stress-Associated Protein 7 (SAP7), an AN1-type zinc-finger protein containing an N-terminal A20 domain and a C-terminal AN1 domain (InterPro: IPR000058 and IPR002653). The PC13 peptide resides between these two domains (**Fig. 4A**). ZmSAP7 localizes to the nucleus and cytoplasm in maize protoplasts and *N. benthamiana* (Lu et al., 2026). SAP proteins have been implicated in early stress responses (Jeffares et al., 2008; Lu et al., 2026). Previous phylogenetic analysis showed that ZmSAP7 clusters with ZmSAP6, ZmSAP10, and *A. thaliana* SAP2 (AtSAP2) within group III SAP proteins (Fu et al., 2022). Notably, the PC13 sequence was conserved only within this SAP subgroup in maize and showed only limited similarity to AtSAP2 (**Fig. 4A**). Comparative analysis of twelve SAP proteins from five Poaceae species (**Fig. 4B**) further revealed that the PC13 motif is highly conserved within the Poaceae species analyzed (**Fig. 4C**), however, no detectable conservation of the PC13 sequence could be observed in non-Poaceae sequences.

**Figure 4:**
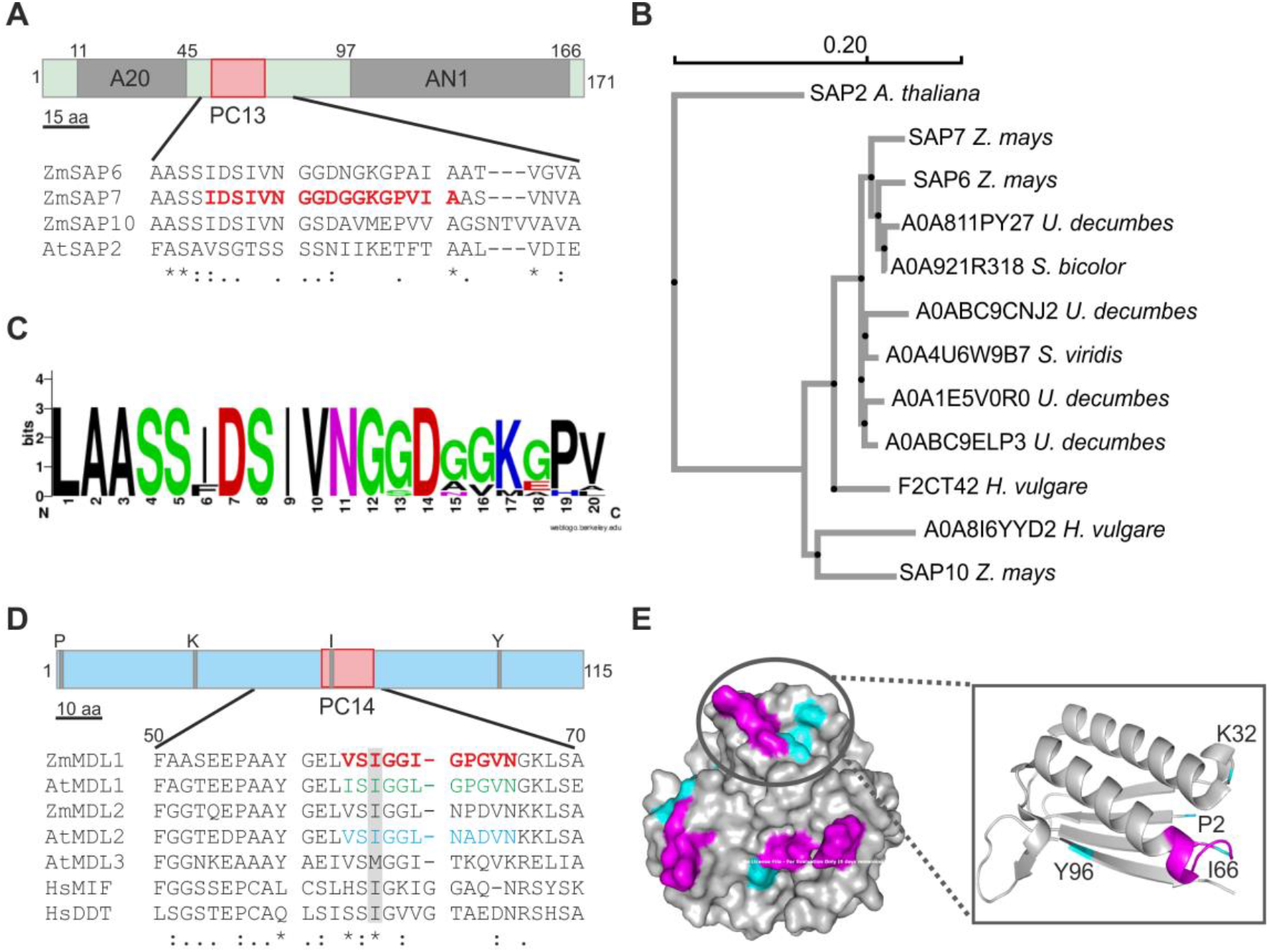
Evolutionary conservation of the PC13 and PC14 propeptides. (A) PC13 is derived from ZmSAP7, a stress-associated protein containing N-terminal A20 and C-terminal AN1 zinc finger domains. The peptide is located in the linker region between the two domains. (B) Phylogenetic tree of 12 protein sequences from different poaceae species (*Zea mays*, *Urochloa decumbens, Setaria viridis, Sorghum bicolor* and *Hordeum vulgare*) identified by BLAST using the PC13 sequence as the query. *Arabidopsis thaliana* SAP2, which lacks the PC13 sequence but belongs to the same phylogenetic group, was used to root the tree (Su *et al*., 2022). (C) Sequence logo illustrating conservation of the PC13 motif across poaceae. (D) PC14 is derived from a ZmMDL (MIF/DDT-like) protein highly conserved across kingdoms and related to the human MIF (HsMIF) and DDT-like (HsDDT) proteins. Sequence comparison of the PC14 region of three *A. thaliana* MDLs (AtMDL1, AtMDL2 and AtMDL3) and two maize ZmMDL orthologues. The conserved isoleucine residue of the tautomerase active site is located within the PC14 motif (grey box). (E) Structural model of the AtMDL1 tetramer (PDB: 8DQ6) showing the surface localization of PC14 (pink) and the tautomerase active site (blue). Inset: AtMDL1 monomer highlighting the putative tautomerase active site residues P2, K32, Y36 (blue), and I66 (pink), which is located within the PC14 motif between α-helices and β-sheets.

PC14 (VSIGGIGPGVN) is derived from the MIF/DDT-like (MDL) protein family. Plant MDLs are homologous to the human migration inhibitory factor (MIF) and DDT-like proteins and are highly conserved across plants, animals, fungi, and protists (Panstruga et al., 2015). Maize contains two PC14-containing MDL homologs closely related to AtMDL1 and AtMDL2, whereas AtMDL3 appears to be dicot-specific (Panstruga et al., 2015). Similar to SAP7, MDL1/2 proteins in wheat, and *Arabidopsis* displayed a nucleo-cytoplasmic localization (Gruner et al., 2021; Zhao et al., 2021). Structural modeling revealed that the conserved PC14 motif encompasses the catalytic isoleucine residue of the MIF-related tautomerase active site (InterPro: IPR001398 and IPR014347; **Fig. 4D, E**). Although plant MDLs lack detectable tautomerase activity, they retain the highly conserved MIF-like fold and assemble into oligomeric complexes (Spiller et al., 2023). Mapping the PC14 sequence onto the *Arabidopsis* MDL1 tetramer (PDB: 8DQ6) localized the peptide to the protein surface at the interface between α-helices and β-sheets (**Fig. 4E**), suggesting that proteolytic excision of PC14 could influence protein complex structure and function.

In summary, PC13 and PC14 represent cryptic phytocytokines derived from functional precursor proteins, ZmSAP7 and ZmMDL1/2, respectively. Their ability to activate immune responses suggests that these proteins possess dual biological functions: acting as intracellular regulators in their precursor form while serving as sources of bioactive extracellular signaling peptides upon proteolytic processing. The evolutionary conservation of the PC13 and PC14 sequences suggests that these regions have been maintained within their precursor proteins, highlighting their potential functional relevance.

### SA-induced apoplastic peptide accumulation coincides with early activation of serine hydrolases

The identification of SA-induced apoplastic peptides raised the question of how these signaling molecules are generated. Phytocytokine production typically requires regulated proteolytic processing, suggesting that activation of extracellular proteases may represent an early step in peptide release. Unlike the previously characterized Zip1 peptide, which is associated with a later activation of apoplastic PLCPs (Koenig et al., 2026; Ziemann et al., 2018), the accumulation of PC13 and PC14 occurs at an early stage 3 h after SA treatment, which suggests the involvement of distinct proteolytic activities. Together with PLCPs, the subtilases (SBTs), a group of serine hydrolases, are among the most prevalent protease classes in the plant apoplast and they have been linked to the production of various phytocytokines. To determine whether serine hydrolases are activated following SA treatment, we performed activity-based protein profiling (ABPP) using the serine hydrolase probe FP-TAMRA, which covalently and irreversibly labels active serine hydrolases. Maize leaves were treated with 2 mM SA or mock solution, and AF was collected at 3, 6, 12, and 24 h after treatment. AF samples were labeled with FP-TAMRA, run on SDS–PAGE, and analyzed by fluorescence scanning. As a specificity control, AF samples were pre-incubated with the SH inhibitors PMSF and DCI. Eight major FP-TAMRA-labelled fluorescent bands were detected across all treatments and time points (**Fig. 5A**, columns A-H). Three bands (**Fig. 5A**, columns E, F, and G) retained residual labeling in the presence of inhibitors, indicating that the inhibitor treatment did not completely block all FP-TAMRA-labelled activities, consistent with previous observations that broad SH inhibitors may not fully inhibit the diverse range of apoplastic hydrolases detected by ABPP (Kaschani et al., 2012). Quantification of the fluorescent bands revealed a 20–40% increase in SH activity at 3 h after SA treatment compared with mock-treated samples (**Fig. 5A**). Signal intensities of bands C–H were significantly increased following SA treatment, indicating enhanced activity of multiple apoplastic serine hydrolases. Analysis of SH activity over time revealed a transient response pattern, with activity decreasing at 6 h after the early 3 h peak, followed by a second increase at 12 h and a subsequent decline at 24 h (**Fig. 5B**). Together, these data demonstrate that SA treatment dynamically modulates the activity of apoplastic serine hydrolases, with the strongest activation occurring at 3 h. This coincides with the time point at which SA-induced apoplastic peptides displaying *PR* gene-inducing activity were detected (**Fig. 1C**). We therefore hypothesized that SBTs might be involved in the generation of peptides in the apoplast 3 h after SA treatment. To address this possibility, we treated maize leaves with mock or SA in the presence or absence of PMSF, a potent plant inhibitor of SBTs and serine carboxypeptidases (SCPLs) (Paulus et al., 2020). AF were collected 3 h after treatment and the apoplastic peptides were enriched using a 10 kDa ultrafiltration device. The resulting APFs were subjected to native peptidomics analysis to identify peptides generated under these conditions. The addition of PMSF strongly altered the apoplastic pool of peptides in both mock-and SA-treated samples (**Table S6**). Interestingly, PC14 was detected exclusively in SA-treated samples in the absence of PMSF, suggesting that its generation depends on PMSF-sensitive serine protease activity including SBTs and SCPLs (**Fig. 5C-D**). Together, these results indicate that serine hydrolase activity is rapidly enhanced following SA treatment and contributes to the generation of apoplastic peptides, including PC14, within 3 h after induction.

**Figure 5:**
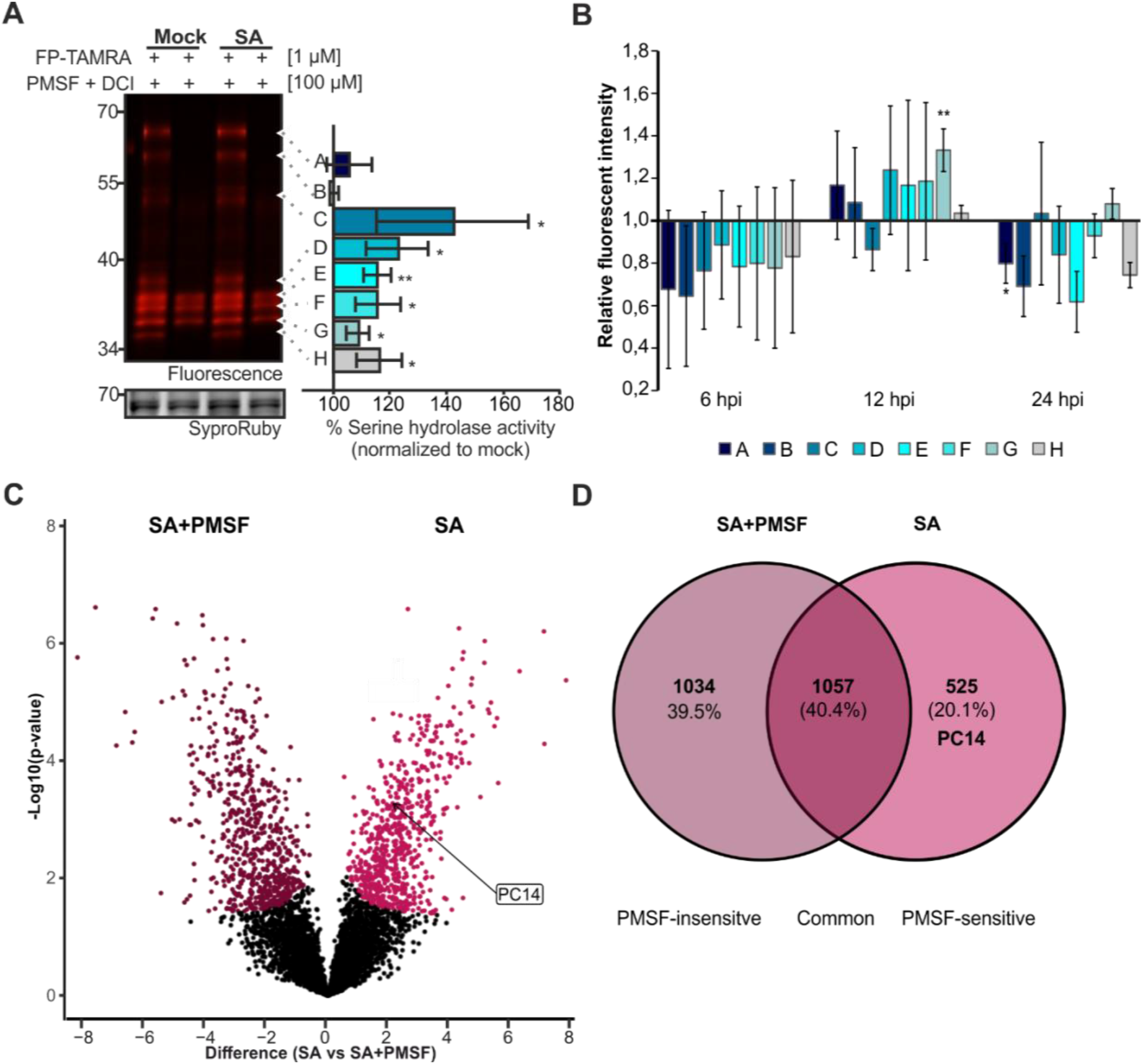
SA treatment rapidly activates apoplastic serine hydrolases. (A) Representative activity-based protein profiling (ABPP) analysis 3 h after treatment. Apoplastic fluid was collected from leaves treated with 2 mM SA or DMSO (mock). Active serine hydrolases (SHs) were labeled with 1 µM FP-TAMRA. Samples pre-incubated with PMSF (100 µM) and DCI (10 µM) served as specificity controls. Labeled proteins were separated by SDS–PAGE and visualized by fluorescence scanning. SyproRuby staining served as a loading control. Fluorescence intensities were normalized to the inhibitor control and the corresponding mock treatment. Data represent three independent biological replicates. Statistical significance was determined using Student’s t-test (P < 0.05, P < 0.01). (B) Apoplastic SH activity after SA treatment over time. ABPP quantification of SH activity of SA labeled samples at 6, 12 and 24 h normalized to mock and the inhibitor control. Plot shows the fluorescence intensity of different bands (A-H). (C) PC14 accumulation depends on PMSF-sensitive serine protease activity. Volcano plot comparing peptide abundance between SA and SA + PMSF treatments. Peptides with significantly altered abundance are highlighted in purple. Peptides enriched in SA-treated samples relative to SA + PMSF represent candidate products of serine protease-mediated processing. (D) Serine protease inhibition alters the apoplastic peptide repertoire. Venn diagram showing peptides uniquely or commonly identified in SA and SA + PMSF-treated samples.

### Cleavage of propeptide ZmMDL1 by apoplastic serine proteases releases PC14

The temporal association between SH activation and PC14 accumulation prompted us to investigate whether apoplastic serine proteases directly contribute to the generation of this phytocytokine. To this end, we fractionated apoplastic fluid collected 3 h after SA treatment by ion-exchange chromatography to enrich for proteolytic activities. A total of 54 fractions were collected, pooled into nine subfractions (A–I), and tested for their ability to cleave recombinant ZmMDL1 (**Fig. 6A**). ZmMDL1 was cloned and fused to a C-terminal His-tag for heterologous production and purification in *E. coli* **(Fig. S2)**. Pool B, containing fractions 7–12, was selected for further analysis because it reproducibly showed ZmMDL1 cleavage activity that was inhibited by PMSF (**Fig. S3**). Within this pool, fraction 7 displayed the strongest ZmMDL1 cleavage activity (**Fig. 6B**). After 60 min incubation with fraction 7, full-length ZmMDL1 was no longer detected, whereas addition of PMSF and DCI prevented cleavage, resulting in the retention of intact ZmMDL1 (**Fig. 6C**). Contrary, ZmMDL1 alone was stable during the course of this experiment. These results show that fraction 7 contains one or more PMSF-sensitive proteolytic activities capable of processing ZmMDL1. Consistent with this observation, ABPP labeling of fraction 7 with FP-TAMRA revealed several active serine hydrolases, which were inhibited by PMSF and DCI, including prominent labeled proteins of approximately 70 kDa (**Fig. 6D**). To determine whether the proteolytic activity present in fraction 7 is capable of generating PC14, mass spectrometry analysis was performed. Purified ZmMDL1 was co-incubated with fraction 7 from SA-treated AF for 60 min in the presence or absence of PMSF. Reaction mixtures were subsequently filtered using a 3 kDa cut-off centrifugal device, and the flow-through fractions were analyzed by LC–MS/MS to identify released peptides. Peptidomic analysis revealed enrichment of one specific peptide corresponding to the PC14 sequence specifically in reactions performed without PMSF (**Fig. 6E**). Additional peptides mapping adjacent to the PC14 region and the N-terminal region of ZmMDL1 were also detected, but at lower abundance (**Table S7**). The identified peptide boundaries suggest that cleavage may occur after the GEL and GKL motifs at the N-and C-terminal regions of PC14, respectively, resulting in the release of the predicted bioactive peptide. Thus, PMSF-sensitive proteolytic activity present in fraction 7 is sufficient to generate PC14-containing peptides from ZmMDL1. Together, these results demonstrate that SA-induced apoplastic serine protease activity contributes to the processing of functional precursor proteins and the generation of bioactive cryptic phytocytokines in maize.

**Figure 6.**
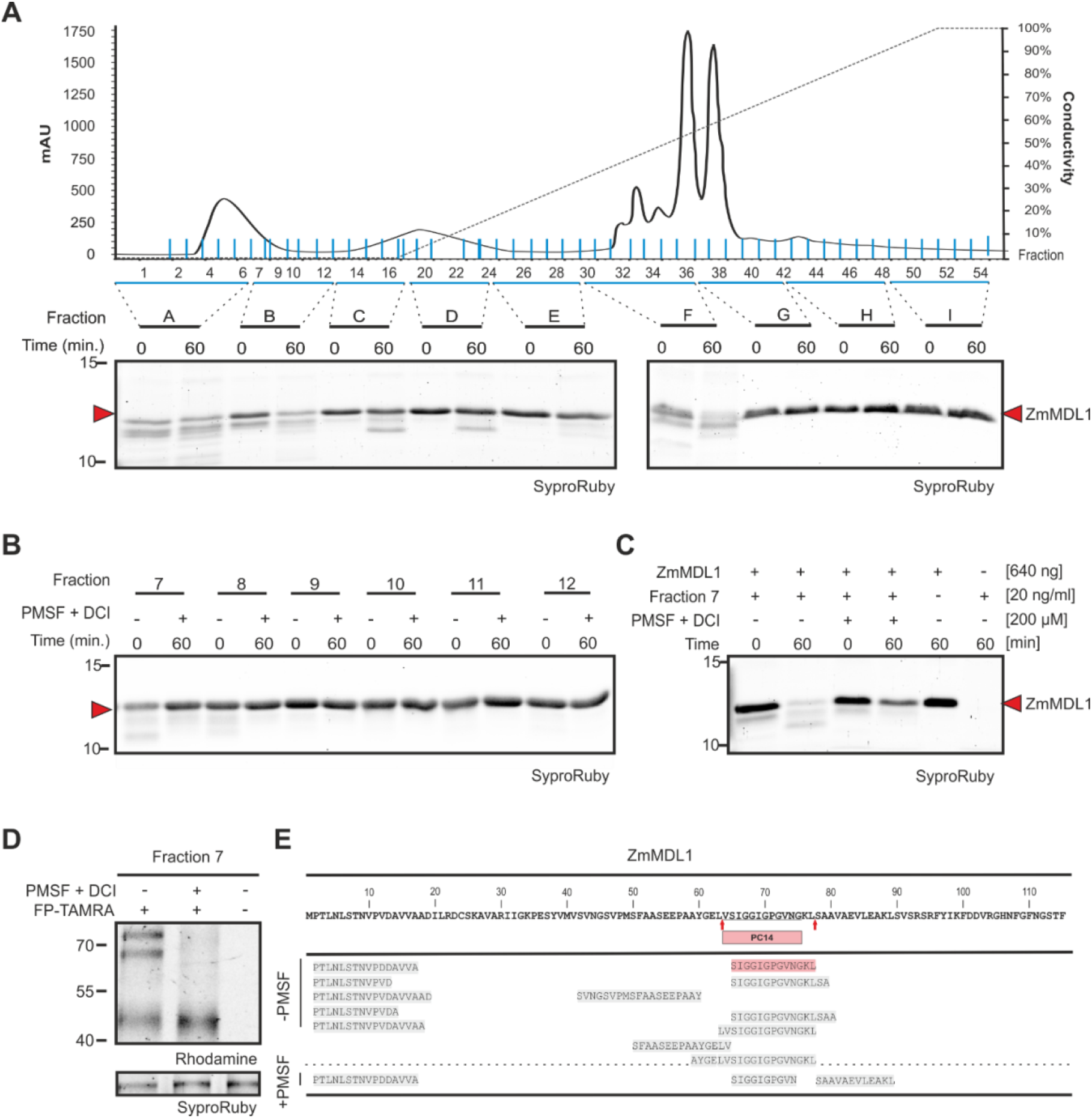
**Apoplastic subtilases mediate PC14 release**. (A) Enrichment of ZmMDL1-cleaving activity. Apoplastic fluid (AF) was collected from maize leaves 3 h after infiltration with 2 mM SA and fractionated by anion-exchange chromatography into 54 fractions. Fractions were grouped (A–I), and ZmMDL1 cleavage activity was assessed for each group. (B) Fraction 7 exhibits the highest ZmMDL1-cleaving activity. Individual fractions (7–12) from group B were analyzed further. Fractions were pre-incubated for 15 min with EDTA (5 mM), pepstatin A (100 µM), and E64 (100 µM), in the presence or absence of PMSF (100 µM) and DCI (100 µM). Proteins were separated by SDS–PAGE, stained with Sypro Ruby, and analyzed by fluorescence scanning. (C) ZmMDL1 cleavage by fraction 7 is PMSF-sensitive. (D) Fraction 7 contains active serine hydrolases. Activity-based protein profiling (ABPP) of Fraction 7 was performed with 1 µM FP-TAMRA following pre-incubation with 100 µM PMSF and 100 µM DCI. Samples were separated by SDS-PAGE and analyzed by fluorescence scanning (E) Fraction 7 releases PC14 from ZmMDL1. Fraction 7 was co-incubated with ZmMDL1 in the presence or absence of PMSF. Samples were filtered using a 3 kDa ultrafiltration device and the flow-through was analyzed by shotgun peptidomics. Shown are all ZmMDL1-His-derived peptides. The peptide highlighted in red was detected in all replicates. Red arrows indicate putative cleavage sites.

## Discussion

This work establishes that SA induces a dynamic release of bioactive peptides into the maize apoplast as early as 3 h after treatment. Peptidomic profiling identified 14 candidate phytocytokines, six of which were validated as inducers of *PR* gene expression. Among these, PC13 and PC14 were demonstrated to have functional activity *in planta*, with opposing effects on virulence of the biotrophic fungal pathogen *U. maydis*. Complementary with the previously characterized phytocytokine Zip1 (Koenig et al., 2026; Ziemann et al., 2018), these findings expand the repertoire of known phytocytokines in maize and reveal a previously unrecognized layer of complexity, where individual peptides can either promote resistance or susceptibility depending on their specific interactions with host signaling networks **(Fig. 7)**. Processing and release of phytocytokines is thereby catalyzed by a spatio-temporal network of proteases. PLCPs were identified first as key players in regulation of apoplastic signaling; however, activity of these proteases was observed in a rather late response 24 h after SA treatment (Misas-Villamil et al., 2016; van der Linde et al., 2012). Metacaspases have been found to confer intracellular processing and release of the Zip1 phytocytokine prior to its maturation and clearance by the PLCPs (Koenig et al., 2026). In this study, we observed an early activation of apoplastic serine hydrolases already three hours after SA treatment and this appears to be required for the apoplastic release of PC14 **(Fig. 7)**. Of particular interest is the fact that both the *in vivo* data from maize apoplastic fluid and the *in vitro* biochemical analysis consistently show that the release of mature PC14 depends on serine protease activity. This activity, in particularly in the case of SBTs, is tightly regulated by subcellular localization and environmental conditions, with pathogen perception and tissue damage-induced apoplastic alkalization promoting SBT activation and peptide precursor processing (Gust et al., 2017; León et al., 2001; Newman et al., 2013; Zhou et al., 2020). Several well-characterized phytocytokines and developmental peptides require SBT-mediated processing for their maturation, highlighting the importance of these proteases in generating bioactive signaling molecules. Thus, our collective observations suggest that at least three classes of proteases (PLCPs, MCs, serine proteases) are involved in the generation of maize phytocytokines in response to just one stimulus, SA **(Fig. 7)**.

**Figure 7.**
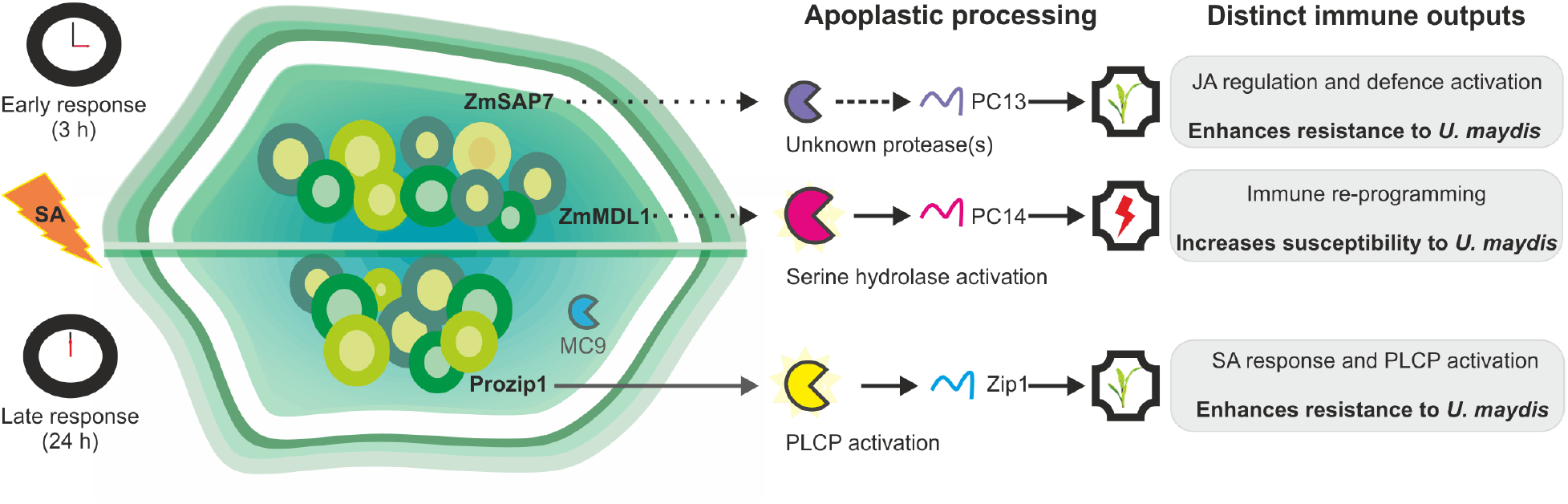
Working model of SA-induced phytocytokines in the maize apoplast. SA triggers an early, transient activation of apoplastic serine hydrolases coinciding with the accumulation of PC13 and PC14. PC13 and PC14 are cryptic phytocytokines derived from the functional proteins ZmSAP7 and ZmMDL1, respectively. Apoplastic serine protease activity contributes to PC14 release. Despite being generated in response to the same stimulus, PC13 and PC14 activate distinct immune programs and have opposing effects on *U. maydis* infection. At later stages, SA-induced PLCP activity promotes the release and clearance of the previously characterized phytocytokine Zip1 from Prozip1, which enhances SA responses and triggers PLCP activation. Solid arrows indicate experimentally supported relationships, whereas dashed arrows represent proposed associations. This model proposes that temporally distinct apoplastic proteolytic activities generate different peptide signals that diversify immune outcomes.

Most phytocytokines identified across different plant families are derived from dedicated precursor proteins with no known or unclear biological function beyond the production of signaling peptides. However, both SAP7 and MDL1, the PC13 and PC14 precursor proteins, have been reported to serve important cellular functions themselves. In animals, SAP proteins regulate innate and adaptive immunity by negatively controlling NF-κB, a key transcriptional regulator of immune responses, inflammation, development, cell proliferation, and survival (Vij & Tyagi, 2006). In plants, SAP proteins are multifunctional regulators involved in ubiquitination, redox homeostasis, growth, development, hormone signaling, and predominantly abiotic stress responses, although they also contribute to plant immunity (Shukla et al., 2021; Vij & Tyagi, 2008). A20/AN1 zinc-finger proteins are conserved across protists, fungi, plants, and animals, exhibiting diverse domain architectures (Vij & Tyagi, 2008). In both plants and mammals, AN1-type proteins localize mainly to the nucleus (Fu et al., 2022).

Three MDL proteins have been identified in Arabidopsis. AtMDL1 and AtMDL2 are constitutively expressed, whereas AtMDL3 shows low basal expression but is strongly induced by biotic and abiotic stress (Panstruga et al., 2015). Arabidopsis MDLs have been linked to flowering time and innate immunity, although their molecular functions remain largely unclear (Gruner et al., 2021). In mammals, MIF is a pleiotropic cytokine regulating inflammation, immunity, tissue repair, cell proliferation, microbial clearance, chemotaxis, cell death, tumorigenesis, and wound healing through intracellular and receptor-mediated extracellular signaling (Sumaiya et al., 2022). In the fungal pathogen *Magnaporthe oryzae*, MIF promotes biotrophic growth by suppressing host cell death (Galli et al., 2023). Despite their evolutionary conservation and extensive characterization in mammals, plant MDL proteins remain poorly understood. Our identification of PC14 as an immune-active peptide derived from the conserved MDL protein raises the possibility that apoplastic proteolytic processing of MDL proteins provides an additional layer of regulation of their biological activity. Whether PC14 function is conserved across MDL proteins remains an open question, given the high conservation of the PC14 sequence among MDL1/2 orthologues. MDL proteins are evolutionarily related to mammalian MIF, which can function both intracellularly and as a secreted cytokine (Kapurniotu et al., 2019; Kleemann et al., 2000). Whether plant MDLs similarly combine intracellular functions with extracellular signaling through proteolytic peptide release remains to be determined.

The identification of non-canonical signaling peptides in the maize apoplast during the early stages of SA treatment highlights an unexpected level of spatial and temporal complexity in phytocytokine regulation. Peptide abundance displayed distinct waves, with a pronounced peak at 3 h, a decline at 6 h, and renewed accumulation at later time points.

These dynamics indicate that the extracellular peptidome is actively remodeled throughout innate immune activation, likely influencing the behavior of cells distal to the infection site. Peptides are well-established regulators of cell fate that control developmental processes in a structure-and concentration-dependent manner (Khavinson et al., 2020), restrict cell death during vascular development and stress, as demonstrated for Kratos (Escamez et al., 2019), and promote tissue regeneration after wounding, as shown for REGENERATION FACTOR1 (REF1) (Yang et al., 2024).

The peptide waves observed in this study suggest that distinct phytocytokines may function at different stages of immunity: early peptides could rapidly amplify defense signaling before transcriptional reprogramming is established, whereas later peptides may sustain immune responses or promote recovery once the initial defense phase subsides. Following this hypothesis, it will be important to investigate the potential mechanistic links between the early release of the phytocytokines PC13 and PC14, which exert opposing effects on immunity, and the subsequent release of later-acting phytocytokines such as Zip1 **(Fig. 7**).

The functional divergence between PC13 and PC14 is one of the key findings of this study. Although both peptides accumulate following SA treatment and induce PR gene expression, they had opposing effects when their activities were tested during *U. maydis* infection. Experimental delivery of PC13 using the Trojan horse system restricted fungal virulence, whereas delivery of PC14 enhanced tumor formation. Thus, immune marker induction alone did not predict the effect of a phytocytokine on disease outcome. Rather, phytocytokines generated in response to the same immune stimulus can exert distinct, even opposing, effects on the plant–pathogen interaction. Such uncoupling of immune activation from disease outcome is increasingly recognized, as pathogens frequently exploit host defense pathways to promote infection. A prominent example is the RALF– FERONIA signaling module, where microbial RALF mimics host FERONIA to facilitate virulence in fungi and nematodes (Masachis et al., 2016; Y. Wang et al., 2024; Zhang et al., 2020). Although the underlying mechanism remains unknown, PC14 may also function as an endogenous susceptibility factor exploited by *U. maydis*. Previous studies have implicated MDL1 homologs, the precursors of PC14, in suppressing programmed cell death (Menuet et al., 2026; Zhao et al., 2021). Consistent with these findings, our results show that the processed PC14 peptide alone is sufficient to promote susceptibility to *U. maydis*. Given that biotrophic pathogens require living host tissue, this activity could contribute to conditions favorable for fungal growth by limiting host cell death.

Transcriptome analysis further associated PC14 with SA signaling. PC14 induces multiple SARD1 and SARD1-like transcription factors, which in Arabidopsis activate ICS genes required for pathogen-induced SA biosynthesis, suggesting that PC14 may reinforce SA signaling through a positive feedback loop. Although this appears inconsistent with the increased susceptibility caused by PC14, *U. maydis* deploys multiple effectors that suppress SA-mediated immunity, including *Um*Cmu1 and *Um*P1p1 (Djamei et al., 2011; Doehlemann et al., 2009; Hemetsberger et al., 2012). Thus, activation of SA-responsive genes is likely outweighed by effector-mediated immune suppression and the ability of PC14 to maintain host cell viability. In contrast, PC13 induced the central SA regulator *NPR1* together with several *TIFY/JAZ* transcription factors, indicating activation of SA signaling while repressing jasmonic acid (JA) responses, consistent with the antagonism between SA and JA pathways during defense against biotrophic pathogens (Ghorbel et al., 2021; Pieterse et al., 2012; Wasternack & Song, 2016; Wu et al., 2012). PC13 also up-regulated genes associated with “programmed cell death” (PCD) and the hypersensitive response, including four membrane-attached complex/perforin (*MACP*) proteins and *LAZ1*, both linked to SA signaling and cell death regulation (Chen et al., 2024; Fu et al., 2022). Together, these results suggest that PC13 enhances resistance by promoting SA-dependent defense and PCD, whereas PC14 favors susceptibility by preserving host cell viability despite activating components of the SA pathway.

Collectively, this study underscores the sophisticated evolutionary strategies by which plants generate defense peptides while pathogens have evolved mechanisms to exploit endogenous signaling molecules for their own benefit. Further investigation into the spatial and temporal regulation involving phytocytokines will further elucidate the precise molecular mechanisms governing these interactions and enhance the understanding of peptide-mediated immunity in plant-microbe systems.

## Materials and Methods

### Plant treatments

Maize plants (*Zea mays,* cv. B73 or KY21) were grown in the phytochamber (28 °C, 15h photoperiod, 40% humidity). For salicylic acid (SA) and peptide treatment, the second leaves of 8-day-old plants were used across independent biological replicates. For apoplastic fluid extraction, detached second leaves from the 8-day-old maize seedlings were vacuum infiltrated with either 2 mM salicylic acid in 1% DMSO or 1% DMSO as a control. Vacuum infiltration was performed in five repetitions of 240 mbar for 10 minutes, each followed by 2 minutes of atmospheric pressure. After infiltration, leaves were incubated at room temperature in closed petri dishes on moist paper towels. At the indicated time points, leaves were submerged in deionized water and vacuum infiltrated in four repetitions of 60 mbar for 15 minutes, each followed by 2 minutes at atmospheric pressure. Leaves were the then gently dried with tissue paper and up to 30 leaves placed cut side down into a 24 ml syringe. The syringe was placed into a 50 ml falcon tube, and everything was centrifuged for 10 min with 2000 x *g* at 4°C. The extracted apoplastic fluid was pooled and transferred into 2 ml tubes. To remove residual leaf debris, the tubes were centrifuged at 10,000 x *g* for 5 min at 4°C. The supernatant was used for further experiments.

### Identification of *Z. mays* immune signalling peptides and protein precursors

To identify maize signalling peptide candidates by mass spectrometry, apoplastic fluid was extracted from the leaves of SA-or mock-treated plants. The apoplastic fluids were filtered using Vivaspin® 10 kDa MWCO Polyethersulfone centrifugal filter units (Sartorius) by centrifugation at 3,000 g for 1 h, until only the dead volume was reached. The resulting filtrate, containing peptides smaller than 10 kDa was frozen and submitted to the CECAD/ZMMK Proteomics Facility of the University of Cologne for MS-based peptidomics analysis. Prior to mass spectrometry, peptide samples were desalted and concentrated using StageTips by the Proteomics Facility.

### Generation of *Ustilago maydis* phytocytokine overexpression strains

The coding sequences of the phytocytokine candidates PC1 to 14 were cloned into the p123 vector system using the Gibson assembly method (Gibson et al., 2009), using the listed primers **(Table S8)**. The vector contains a constitutive promoter (pro*^pit2^)* for overexpression, a terminator (*Tnos*), and a resistance carboxin resistance (*cbx*) cassette for selection. The assembled plasmids were transformed into TOP10 *E. coli* cells by heat shock for plasmid amplification. Purified plasmids were linearized by *SspI* restriction enzyme to allow the homologous recombination into the *ip* locus of *U. maydis*. For fungal transformation, 50 μl protoplasts of *U. maydis* (SG200; solo pathogenic mutant) were transfected by adding 5 μg of linearized plasmid, 1 μl Heparin and 0.5 ml STC / 40% PEG (sterile) incubating for 15 minutes on ice. Transformed protoplasts were plated onto regeneration agar prepared as two-layer system, consisting of a top layer without selection and a bottom layer containing carboxin. Plates were incubated at 28°C for 4-5 days. Colonies were picked and singled out onto Potato-Dextrose-agar (PDA) plates for two days at 28°C. Single colonies were inoculated into 2 ml culture and grown overnight for genomic DNA extraction (Hoffman & Winston, 1987). The extracted DNA was digested by HindIII, separated on 0.9% agarose gel (100 V, 2 h), and blotted on a nylon membrane for Southern Blot analysis. Integration at the *ip* locus was confirmed by hybridization with dixogenin (DIG)-labelled *cbx* probe. Successfully confirmed strains were used for subsequent infection assays.

### *Ustilago maydis* cultivation and infection assays

SG200, SG200_*Zm*PC1, SG200_*Zm*PC10, SG200_*Zm*PC11, SG200_*Zm*PC12, SG200_*Zm*PC13 and SG200_*Zm*PC14 were grown in YEPS light medium to an OD600 of 0.8. The cultures were pelleted at 2,850 x *g* for 7 minutes and to remove remaining cultivation media the pellets were washed with *dd*H2O. The washed pallets were resuspended to a final OD600 of 1.0. To verify the filamentation, 10 μl droplets of each culture were placed on charcoal plates. For infection assays, 7-day-old maize seedlings were inoculated by injecting the fungal suspension into the stem using a 1 ml syringe with a needle. The disease symptoms were evaluated at 12 dpi using a standard disease rating scale for *Ustilago maydis* comprising: healthy plants, chlorosis, small tumours (<2 mm), normal tumours (2 to 10 mm diameter), heavy tumours (>10 mm or stunted growth) and dead plants. Plants were categorized based on their phenotype, and disease severity of the overexpression strains was compared with that of the SG200 control strain.

### Expression and purification of ZmMDL1

The coding sequence of *Zm*MDL1 was amplified from *Z. mays* B73 complementary DNA (cDNA) using the oligonucleotides M0-MDL1-Fw and M0-MDL1-Rv **(Table S8)**. Recombinant *Zm*MDL1 protein was heterologously expressed and purified by immobilized metal affinity chromatography (IMAC) using a 1 mL HisTrap FF Crude column (Cytiva) on an ÄKTA Start chromatography system. Prior to sample loading, the column was equilibrated with Ni-NTA binding buffer. Purification was performed using an automated protocol consisting of 5 column volumes (CV) for equilibration, 15 CV for washing, and 7 CV for elution. Eluted proteins were collected in 1 mL fractions using Ni-NTA elution buffer. Fractions containing *Zm*MDL1 were pooled and further purified by size-exclusion chromatography (SEC) on an ÄKTA chromatography system (GE Healthcare Life Sciences, Buckinghamshire, UK) equipped with a Superdex 75 16/600 column. The column was equilibrated with storage buffer containing 50 mM Tris-HCl (pH 7.0) and 150 mM NaCl. Purified protein fractions were used for subsequent experiments.

### Fractionation of Apoplastic Fluid by Anion-Exchange Chromatography

Apoplastic fluid (5 mL) was pre-incubated with 30 µM pepstatin A (PepA) and 30 µM E-64 for 5 min and subsequently passed through a 0.22 µm syringe filter before chromatography. The filtered sample was loaded onto a 1 mL Mono Q anion-exchange column (GE Healthcare) equilibrated with 20 mM sodium phosphate buffer (pH 6.0). The column was washed with 2 mL of equilibration buffer, and the flow-through and wash fractions were collected in 1 mL aliquots. Bound proteins were eluted with a 20 mL linear gradient of 0–1 M NaCl in 20 mM sodium phosphate buffer (pH 6.0), and 500 µL fractions were collected throughout the gradient. The column was subsequently washed with 5 mL of 1 M NaCl in 20 mM sodium phosphate buffer (pH 6.0), and the eluate was collected in 1 mL fractions.

### Activity-based protein profiling of serine hydrolases

Serine hydrolase activity was analyzed by activity-based protein profiling (ABPP) using the fluorophosphonate-based probe FP-TAMRA. Total protein extract (TE), apoplastic fluid (AF), or fractionated apoplastic fluid was incubated in darkness for 2 h at room temperature in 20 mM sodium acetate (pH 6.0), DMSO and 1 µM of the FP-TAMRA probe. Prior to the labelling, one set of samples were pre-incubated with 100 µM of the inhibitors PMSF and DCI for 15 minutes at room temperature as negative control. Labeling was terminated by the addition of 1x SDS-loading dye. Samples were heated to 95°C for 5 min and separated on 12% SDS–PAGE in the dark. FP-TAMRA-labeled proteins were detected using a ChemiDoc imaging system (Bio-Rad, Hercules, CA, USA) with the rhodamine filter settings (excitation: 532 nm, emission: 580 nm). To visualize total protein content, gels were stained overnight with SYPRO Ruby (Thermo Fisher Scientific, S12000) according to the manufacturer’s protocol. Fluorescence and total protein signals were quantified using Image Lab software (Bio-Rad, Hercules, CA, USA).

### RNA sequencing and transcriptome analysis

For RNA-seq samples, the second leaves of the 8-day-old maize seedlings were infiltered with 4 µM chemically synthesized peptides using needleless syringes. The syringe was applied to the bottom side of the leaves and the solution was infiltrated by gentle pressure. Infiltration was conducted using four points, comprising two upper and two lower points. The section between the two points was utilized for the analysis of gene expression and frozen in liquid nitrogen. Total RNA was extracted from maize leaves harvested 2, 6, 12, and 24 h after treatment with MAP1, Zip1, PC13, or PC14 by using TRIzol (Thermo Fisher, Waltham, USA) according to the manufacturer’s protocol. Three independent biological replicates were prepared for each treatment and time point. Library preparation and sequencing of the RNA samples was performed by Novogene (UK/Munich). RNA libraries were generated using the Illumina TruSeq Stranded mRNA Library Preparation Kit (Illumina, San Diego, CA, USA) and sequenced as paired-end reads on an Illumina NovaSeq X Plus platform. Raw sequencing reads were subjected to quality control and filtering using standard parameters. Filtered reads were aligned to the *Zea mays* B73 reference genome version 5 (MaizeGDB). Read mapping was performed using HISAT2, and gene-level read counts were used for differential gene expression analysis with the DESeq2 package in R. Gene expression levels were calculated as counts per million (CPM), and transcripts per million (TPM) values were determined for expression normalization. In addition, a standard RNA-seq bioinformatics analysis pipeline was performed by Novogene.

## Acknowledgements

We are grateful to all our team members and colleagues who contributed to work on this project. We thank Jan-Wilm Lackmann and the Proteomics core facility Cologne for performing peptidomics and related data analysis. We acknowledge Jan Muelhoefer and Ute Meyer for their excellent technical support. This work was supported by the Deutsche Forschungsgemeinschaft (DFG) under project DO 1421/5-2, the SFB1403 (project no. 414786233), and the Cluster of Excellence on Plant Sciences (CEPLAS, EXC 2048/1— project ID: 390686111).

## Data availability

RNA-seq raw data are publicly available on the NCBI Gene Expression Omnibus (accession number GSE343547). Mass Spectrometry data are publicly accessible in the PRIDE database (accession number: PXD083485). Biological material mentioned in the manuscript can be requested from the corresponding authors (GD, JM) upon reasonable request.

## Author Contributions

PK, HL, JCMV, and GD designed the study and the experiments, PK and HL conducted the experimental work. PK, HL and ZZ performed peptidomics and RNAseq data analysis. PK, HL, ZZ, JCMV, GD wrote the manuscript with contributions from all authors.

## Supplementary Information

**Figure S1.** Peptide abundance over time after SA treatment.

**Figure S2.** MIF-His protein purification

**Figure S3.** Stability of MIF in different fractions in the presence of PMSF.

**Table S1.** Peptides identified by MS-based peptidomics analysis of maize leaf apoplastic fluid collected 3 h after salicylic acid (SA) or buffer (mock) infiltration

**Table S2.** Peptides identified by MS-based peptidomics analysis of maize leaf apoplastic fluid collected 6 h, 9 h and 12 h after salicylic acid (SA) or buffer (mock) infiltration

**Table S3.** Selected peptides from MS analysis of APF

**Table S4.** Table of differential gene expression analysis.

**Table S5.** Significantly enriched GO terms for Biological Processes (BP) in the upregulated genes at 2 hpi and 24 hpi.

**Table S6.** Peptides identified by MS-based peptidomics analysis of maize leaf apoplastic fluid collected 3h following SA, SA+PMSF or buffer (mock) infiltration

**Table S7.** Cleavage products of ZmMDL1-His (Zm00001eb226780) following co-incubation with fraction 7

**Table S8.** Oligonucleotides

## References

Amano, Y., Tsubouchi, H., Shinohara, H., Ogawa, M., & Matsubayashi, Y. (2007). Tyrosine-sulfated glycopeptide involved in cellular proliferation and expansion in *Arabidopsis*. Proceedings of the National Academy of Sciences, 104(46), 18333–18338. 10.1073/pnas.0706403104

Beloshistov, R. E., Dreizler, K., Galiullina, R. A., Tuzhikov, A. I., Serebryakova, M. V., Reichardt, S., Shaw, J., Taliansky, M. E., Pfannstiel, J., Chichkova, N. V., Stintzi, A., Schaller, A., & Vartapetian, A. B. (2018). Phytaspase-mediated precursor processing and maturation of the wound hormone systemin. New Phytologist, 218(3), 1167–1178. 10.1111/nph.14568

Boller, T., & Felix, G. (2009). A Renaissance of Elicitors: Perception of Microbe-Associated Molecular Patterns and Danger Signals by Pattern-Recognition Receptors. Annual Review of Plant Biology, 60(1), 379–406. 10.1146/annurev.arplant.57.032905.105346

Butenko, M. A. (2003). INFLORESCENCE DEFICIENT IN ABSCISSION Controls Floral Organ Abscission in Arabidopsis and Identifies a Novel Family of Putative Ligands in Plants. THE PLANT CELL ONLINE, 15(10), 2296–2307. 10.1105/tpc.014365

Castaldi, V., Wicaksono, W. A., Criscuolo, M. C., Gualtieri, L., Langella, E., Lelio, I. D., Monti, S. M., Filippis, F. D., Berg, G., & Rao, R. (2026). Prosystemin-derived signals: Bridging leaf microbiome dynamics and defense activation. Environmental Microbiome, 21(1), 71. 10.1186/s40793-026-00885-9

Chen, C., Buscaill, P., Sanguankiattichai, N., Huang, J., Kaschani, F., Kaiser, M., & Hoorn, R. A. L. van der. (2024). Extracellular plant subtilases dampen cold-shock peptide elicitor levels. Nature Plants, 10(11), 1749–1760. 10.1038/s41477-024-01815-8

Choi, H. W., & Klessig, D. F. (2016). DAMPs, MAMPs, and NAMPs in plant innate immunity. BMC Plant Biology, 16(1), 232. 10.1186/s12870-016-0921-2

Depotter, J. R. L., Villamil, J. C. M., & Doehlemann, G. (2022, May). Maize immune signalling peptide ZIP1 evolved de novo from a retrotransposon. 10.1101/2022.05.18.492421

Djamei, A., Schipper, K., Rabe, F., Ghosh, A., Vincon, V., Kahnt, J., Osorio, S., Tohge, T., Fernie, A. R., Feussner, I., Feussner, K., Meinicke, P., Stierhof, Y.-D., Schwarz, H., Macek, B., Mann, M., & Kahmann, R. (2011). Metabolic priming by a secreted fungal effector. Nature, 478(7369), 395–398. 10.1038/nature10454

Doehlemann, G., Linde, K. van der, Aßmann, D., Schwammbach, D., Hof, A., Mohanty, A., Jackson, D., & Kahmann, R. (2009). Pep1, a Secreted Effector Protein of Ustilago maydis, Is Required for Successful Invasion of Plant Cells. PLoS Pathogens, 5(2), e1000290. 10.1371/journal.ppat.1000290

Escamez, S., Stael, S., Vainonen, J. P., Willems, P., Jin, H., Kimura, S., van Breusegem, F., Gevaert, K., Wrzaczek, M., & Tuominen, H. (2019). Extracellular peptide Kratos restricts cell death during vascular development and stress in Arabidopsis. Journal of Experimental Botany, 70(7), 2199–2210. 10.1093/jxb/erz021

Fu, Q., Duan, H., Cao, Y., Li, Y., Lin, X., Pang, H., Yang, Q., Li, W., Fu, F., Zhang, Y., & Yu, H. (2022). Comprehensive Identification and Functional Analysis of Stress-Associated Protein (SAP) Genes in Osmotic Stress in Maize. International Journal of Molecular Sciences, 23(22), 14010. 10.3390/ijms232214010

Galli, M., Jacob, S., Zheng, Y., Ghezellou, P., Gand, M., Albuquerque, W., Imani, J., Allasia, V., Coustau, C., Spengler, B., Keller, H., Thines, E., & Kogel, K.-H. (2023). MIF-like domain containing protein orchestrates cellular differentiation and virulence in the fungal pathogen Magnaporthe oryzae. iScience, 26(9), 107565. 10.1016/j.isci.2023.107565

Ghorbel, M., Brini, F., Sharma, A., & Landi, M. (2021). Role of jasmonic acid in plants: The molecular point of view. Plant Cell Reports, 40(8), 1471–1494. 10.1007/s00299-021-02687-4

Gibson, D. G., Young, L., Chuang, R.-Y., Venter, J. C., Hutchison, C. A., & Smith, H. O. (2009). Enzymatic assembly of DNA molecules up to several hundred kilobases. Nature Methods, 6(5), 343–345. 10.1038/nmeth.1318

Gruner, K., Leissing, F., Sinitski, D., Thieron, H., Axstmann, C., Baumgarten, K., Reinstädler, A., Winkler, P., Altmann, M., Flatley, A., Jaouannet, M., Zienkiewicz, K., Feussner, I., Keller, H., Coustau, C., Falter-Braun, P., Feederle, R., Bernhagen, J., & Panstruga, R. (2021). Chemokine-like MDL proteins modulate flowering time and innate immunity in plants. Journal of Biological Chemistry, 296, 100611. 10.1016/j.jbc.2021.100611

Gust, A. A., Pruitt, R., & Nürnberger, T. (2017). Sensing Danger: Key to Activating Plant Immunity. Trends in Plant Science, 22(9), 779–791. 10.1016/j.tplants.2017.07.005

Hander, T., Fernández-Fernández, Á. D., Kumpf, R. P., Willems, P., Schatowitz, H., Rombaut, D., Staes, A., Nolf, J., Pottie, R., Yao, P., Gonçalves, A., Pavie, B., Boller, T., Gevaert, K., Breusegem, F. V., Bartels, S., & Stael, S. (2019). Damage on plants activates Ca ^2+^ - dependent metacaspases for release of immunomodulatory peptides. Science, 363(6433). 10.1126/science.aar7486

Hemetsberger, C., Herrberger, C., Zechmann, B., Hillmer, M., & Doehlemann, G. (2012). The Ustilago maydis Effector Pep1 Suppresses Plant Immunity by Inhibition of Host Peroxidase Activity. PLoS Pathogens, 8(5), e1002684. 10.1371/journal.ppat.1002684

Hou, S., Liu, D., Huang, S., Luo, D., Liu, Z., Xiang, Q., Wang, P., Mu, R., Han, Z., Chen, S., Chai, J., Shan, L., & He, P. (2021). The Arabidopsis MIK2 receptor elicits immunity by sensing a conserved signature from phytocytokines and microbes. Nature Communications, 12(1), 5494. 10.1038/s41467-021-25580-w

Hou, S., Wang, X., Chen, D., Yang, X., Wang, M., Turrà, D., Pietro, A. D., & Zhang, W. (2014). The Secreted Peptide PIP1 Amplifies Immunity through Receptor-Like Kinase 7. PLoS Pathogens, 10(9), e1004331. 10.1371/journal.ppat.1004331

Jeffares, D. C., Penkett, C. J., & Bähler, J. (2008). Rapidly regulated genes are intron poor. Trends in Genetics, 24(8), 375–378. 10.1016/j.tig.2008.05.006

Kapurniotu, A., Gokce, O., & Bernhagen, J. (2019). The Multitasking Potential of Alarmins and Atypical Chemokines. Frontiers in Medicine, 6, 3. 10.3389/fmed.2019.00003

Kaschani, F., Nickel, S., Pandey, B., Cravatt, B. F., Kaiser, M., & Hoorn, R. A. L. van der. (2012). Selective inhibition of plant serine hydrolases by agrochemicals revealed by competitive ABPP. Bioorganic & Medicinal Chemistry, 20(2), 597–600. 10.1016/j.bmc.2011.06.040

Khavinson, V., Linkova, N., Diatlova, A., & Trofimova, S. (2020). Peptide Regulation of Cell Differentiation. Stem Cell Reviews and Reports, 16(1), 118–125. 10.1007/s12015-019-09938-8

Kleemann, R., Hausser, A., Geiger, G., Mischke, R., Burger-Kentischer, A., Flieger, O., Johannes, F.-J., Roger, T., Calandra, T., Kapurniotu, A., Grell, M., Finkelmeier, D., Brunner, H., & Bernhagen, J. (2000). Intracellular action of the cytokine MIF to modulate AP-1 activity and the cell cycle through Jab1. Nature, 408(6809), 211–216. 10.1038/35041591

Koenig, M., Moser, D., Leusner, J., Depotter, J. R. L., Doehlemann, G., & Villamil, J. M. (2023). Maize Phytocytokines Modulate Pro-Survival Host Responses and Pathogen Resistance. Molecular Plant-Microbe Interactions®, 36(9), 592–604. 10.1094/MPMI-01-23-0005-R

Koenig, M., Sorger, Z., Kakanj, P., Dewes, P., Mantz, M., Perrar, A., Sivaramakrishnan, M., Stael, S., Chandrasekar, B., Huesgen, P. F., Villamil, J. M., & Doehlemann, G. (2026). Processing and release of the maize phytocytokine Zip1. *Plant Physiology*, kiag533. 10.1093/plphys/kiag533

Lalun, V. O., Breiden, M., Galindo-Trigo, S., Smakowska-Luzan, E., Simon, R. G., & Butenko, M. A. (2024). A dual function of the IDA peptide in regulating cell separation and modulating plant immunity at the molecular level. eLife, 12, RP87912. 10.7554/eLife.87912

Lecourieux, D., Mazars, C., Pauly, N., Ranjeva, R., & Pugin, A. (2002). Analysis and Effects of Cytosolic Free Calcium Increases in Response to Elicitors in *Nicotiana plumbaginifolia* Cells. The Plant Cell, 14(10), 2627–2641. 10.1105/tpc.005579

León, J., Rojo, E., & Sánchez-Serrano, J. J. (2001). Wound signalling in plants. Journal of Experimental Botany, 52(354), 1–9. 10.1093/jexbot/52.354.1

Liu, P., Shi, C., Liu, S., Lei, J., Lu, Q., Hu, H., Ren, Y., Zhang, N., Sun, C., Chen, L., Jiang, Y., Feng, L., Zhang, T., Zhong, K., Liu, J., Zhang, J., Zhang, Z., Sun, B., Chen, J.,…Yang, J. (2024). Author Correction: A papain-like cysteine protease-released small signal peptide confers wheat resistance to wheat yellow mosaic virus. Nature Communications, 15(1), 991. 10.1038/s41467-024-45406-9

Lu, F., Li, Y., Duan, H., He, L., He, R., Tang, Q., Wang, Y., Fu, F., Lu, Y., & Yu, H. (2026). Maize stress-associated proteins ZmSAP1 and ZmSAP7 positively regulate salt stress tolerance. Plant Physiology and Biochemistry, 231, 111028. 10.1016/j.plaphy.2026.111028

Ma, M., Jiang, W., & Zhou, R. (2024). DAMPs and DAMP-sensing receptors in inflammation and diseases. Immunity, 57(4), 752–771. 10.1016/j.immuni.2024.03.002

Masachis, S., Segorbe, D., Turrà, D., Leon-Ruiz, M., Fürst, U., Ghalid, M. E., Leonard, G., López-Berges, M. S., Richards, T. A., Felix, G., & Pietro, A. D. (2016). Correction: Corrigendum: A fungal pathogen secretes plant alkalinizing peptides to increase infection. Nature Microbiology, 1(1), 16073. 10.1038/nmicrobiol.2016.73

Meng, X., & Zhang, S. (2013). MAPK Cascades in Plant Disease Resistance Signaling. Annual Review of Phytopathology, 51(1), 245–266. 10.1146/annurev-phyto-082712-102314

Misas-Villamil, J. C., Hoorn, R. A. L. van der, & Doehlemann, G. (2016). Papain-like cysteine proteases as hubs in plant immunity. New Phytologist, 212(4), 902–907. 10.1111/nph.14117

Narváez-Vásquez, J., Pearce, G., & Ryan, C. A. (2005). The plant cell wall matrix harbors a precursor of defense signaling peptides. Proceedings of the National Academy of Sciences, 102(36), 12974–12977. 10.1073/pnas.0505248102

Newman, M.-A., Sundelin, T., Nielsen, J. T., & Erbs, G. (2013). MAMP (microbe-associated molecular pattern) triggered immunity in plants. Frontiers in Plant Science, 4. 10.3389/fpls.2013.00139

Ngou, B. P. M., Ding, P., & Jones, J. D. G. (2022). Thirty years of resistance: Zig-zag through the plant immune system. The Plant Cell, 34(5), 1447–1478. 10.1093/plcell/koac041

Panstruga, R., Baumgarten, K., & Bernhagen, J. (2015). Phylogeny and evolution of plant macrophage migration inhibitory factor/D-dopachrome tautomerase-like proteins. BMC Evolutionary Biology, 15(1), 64. 10.1186/s12862-015-0337-x

Paulus, J. K., Kourelis, J., Ramasubramanian, S., Homma, F., Godson, A., Hörger, A. C., Hong, T. N., Krahn, D., Carballo, L. O., Wang, S., Win, J., Smoker, M., Kamoun, S., Dong, S., & Hoorn, R. A. L. van der. (2020). Extracellular proteolytic cascade in tomato activates immune protease Rcr3. Proceedings of the National Academy of Sciences, 117(29), 17409–17417. 10.1073/pnas.1921101117

Pearce, G., Moura, D. S., Stratmann, J., & Ryan, C. A. (2001). Production of multiple plant hormones from a single polyprotein precursor. Nature, 411(6839), 817–820. 10.1038/35081107

Pearce, G., Yamaguchi, Y., Barona, G., & Ryan, C. A. (2010a). A subtilisin-like protein from soybean contains an embedded, cryptic signal that activates defense-related genes. Proceedings of the National Academy of Sciences, 107(33), 14921–14925. 10.1073/pnas.1007568107

Pearce, G., Yamaguchi, Y., Barona, G., & Ryan, C. A. (2010b). A subtilisin-like protein from soybean contains an embedded, cryptic signal that activates defense-related genes. Proceedings of the National Academy of Sciences, 107(33), 14921–14925. 10.1073/pnas.1007568107

Peng, Y., van Wersch, R., & Zhang, Y. (2018). Convergent and Divergent Signaling in PAMP-Triggered Immunity and Effector-Triggered Immunity. Molecular Plant-Microbe Interactions®, 31(4), 403–409. 10.1094/MPMI-06-17-0145-CR

Pieterse, C. M. J., Does, D. V. der, Zamioudis, C., Leon-Reyes, A., & Wees, S. C. M. V. (2012). Hormonal Modulation of Plant Immunity. Annual Review of Cell and Developmental Biology, 28(1), 489–521. 10.1146/annurev-cellbio-092910-154055

Reichardt, S., Piepho, H.-P., Stintzi, A., & Schaller, A. (2020). Peptide signaling for drought-induced tomato flower drop. Science, 367(6485), 1482–1485. 10.1126/science.aaz5641

Rodríguez, A., Martín, M., Albasanz, J. L., Barrachina, M., Espinosa, J. C., Torres, J. M., & Ferrer, I. (2006). Group I mGluR signaling in BSE-infected bovine-PrP transgenic mice. Neuroscience Letters, 410(2), 115–120. 10.1016/j.neulet.2006.09.084

Schardon, K., Hohl, M., Graff, L., Pfannstiel, J., Schulze, W., Stintzi, A., & Schaller, A. (2016). Precursor processing for plant peptide hormone maturation by subtilisin-like serine proteinases. Science, 354(6319), 1594–1597. 10.1126/science.aai8550

Schmelz, E. A., Carroll, M. J., LeClere, S., Phipps, S. M., Meredith, J., Chourey, P. S., Alborn, H. T., & Teal, P. E. A. (2006). Fragments of ATP synthase mediate plant perception of insect attack. Proceedings of the National Academy of Sciences, 103(23), 8894–8899. 10.1073/pnas.0602328103

Shen, W., Liu, J., & Li, J.-F. (2019). Type-II Metacaspases Mediate the Processing of Plant Elicitor Peptides in Arabidopsis. Molecular Plant, 12(11), 1524–1533. 10.1016/j.molp.2019.08.003

Shukla, V., Choudhary, P., Rana, S., & Muthamilarasan, M. (2021). Structural evolution and function of stress associated proteins in regulating biotic and abiotic stress responses in plants. Journal of Plant Biochemistry and Biotechnology, 30(4), 779–792. 10.1007/s13562-021-00704-x

Spiller, L., Manjula, R., Leissing, F., Basquin, J., Bourilhon, P., Sinitski, D., Brandhofer, M., Levecque, S., Sabelleck, B., Feederle, R., Flatley, A., Panstruga, R., Bernhagen, J., & Lolis, E. (2023, January). Structures of Arabidopsis thaliana MDL Proteins and Synergistic Effects with the Cytokine MIF on Human Receptors. 10.1101/2023.01.30.525655

Sumaiya, K., Langford, D., Natarajaseenivasan, K., & Shanmughapriya, S. (2022). Macrophage migration inhibitory factor (MIF): A multifaceted cytokine regulated by genetic and physiological strategies. Pharmacology & Therapeutics, 233, 108024. 10.1016/j.pharmthera.2021.108024

van Der Linde, K., Hemetsberger, C., Kastner, C., Kaschani, F., van Der Hoorn, R. A. L., Kumlehn, J., & Doehlemann, G. (2012). A Maize Cystatin Suppresses Host Immunity by Inhibiting Apoplastic Cysteine Proteases. The Plant Cell, 24(3), 1285–1300. 10.1105/tpc.111.093732

van Der Linde, K., Timofejeva, L., Egger, R. L., Ilau, B., Hammond, R., Teng, C., Meyers, B. C., Doehlemann, G., & Walbot, V. (2018). Pathogen Trojan Horse Delivers Bioactive Host Protein to Alter Maize Anther Cell Behavior in Situ. The Plant Cell, 30(3), 528–542. 10.1105/tpc.17.00238

Vij, S., & Tyagi, A. K. (2006). Genome-wide analysis of the stress associated protein (SAP) gene family containing A20/AN1 zinc-finger(s) in rice and their phylogenetic relationship with Arabidopsis. Molecular Genetics and Genomics, 276(6), 565–575. 10.1007/s00438-006-0165-1

Vij, S., & Tyagi, A. K. (2008). A20/AN1 zinc-finger domain-containing proteins in plants and animals represent common elements in stress response. Functional & Integrative Genomics, 8(3), 301–307. 10.1007/s10142-008-0078-7

Wang, S., Zhang, J., Gao, X., Bao, X., Liu, S., Qin, R., Xin, B., Li, P., Zhang, B., & Wu, L. (2026). Plant non-canonical peptides: From identification to mechanisms. Plant Communications, 7(3), 101739. 10.1016/j.xplc.2026.101739

Wang, Y., Liu, X., Yuan, B., Chen, X., Zhao, H., Ali, Q., Zheng, M., Tan, Z., Yao, H., Zheng, S., Wu, J., Xu, J., Shi, J., Wu, H., Gao, X., & Gu, Q. (2024). Fusarium graminearum rapid alkalinization factor peptide negatively regulates plant immunity and cell growth via the FERONIA receptor kinase. Plant Biotechnology Journal, 22(7), 1800–1811. 10.1111/pbi.14303

Wasternack, C., & Song, S. (2016). Jasmonates: Biosynthesis, metabolism, and signaling by proteins activating and repressing transciption. *Journal of Experimental Botany*, erw443. 10.1093/jxb/erw443

Wu, Y., Zhang, D., Chu, J. Y., Boyle, P., Wang, Y., Brindle, I. D., De Luca, V., & Després, C. (2012). The Arabidopsis NPR1 Protein Is a Receptor for the Plant Defense Hormone Salicylic Acid. Cell Reports, 1(6), 639–647. 10.1016/j.celrep.2012.05.008

Yang, W., Zhai, H., Wu, F., Deng, L., Chao, Y., Meng, X., Chen, Q., Liu, C., Bie, X., Sun, C., Yu, Y., Zhang, X., Zhang, X., Chang, Z., Xue, M., Zhao, Y., Meng, X., Li, B., Zhang, X.,…Li, C. (2024). Peptide REF1 is a local wound signal promoting plant regeneration. Cell, 187(12), 3024–3038.e14. 10.1016/j.cell.2024.04.040

Zhang, X., Peng, H., Zhu, S., Xing, J., Li, X., Zhu, Z., Zheng, J., Wang, L., Wang, B., Chen, J., Ming, Z., Yao, K., Jian, J., Luan, S., Coleman-Derr, D., Liao, H., Peng, Y., Peng, D., & Yu, F. (2020). Nematode-Encoded RALF Peptide Mimics Facilitate Parasitism of Plants through the FERONIA Receptor Kinase. Molecular Plant, 13(10), 1434–1454. 10.1016/j.molp.2020.08.014

Zhao, M., Chang, Q., Liu, Y., Sang, P., Kang, Z., & Wang, X. (2021). Functional Characterization of the Wheat Macrophage Migration Inhibitory Factor TaMIF1 in Wheat-Stripe Rust (Puccinia striiformis) Interaction. Biology, 10(9), 878. 10.3390/biology10090878

Zhou, F., Emonet, A., Dénervaud Tendon, V., Marhavy, P., Wu, D., Lahaye, T., & Geldner, N. (2020). Co-incidence of Damage and Microbial Patterns Controls Localized Immune Responses in Roots. Cell, 180(3), 440–453.e18. 10.1016/j.cell.2020.01.013

Ziemann, S., van Der Linde, K., Lahrmann, U., Acar, B., Kaschani, F., Colby, T., Kaiser, M., Ding, Y., Schmelz, E., Huffaker, A., Holton, N., Zipfel, C., & Doehlemann, G. (2018). An apoplastic peptide activates salicylic acid signalling in maize. Nature Plants, 4(3), 172–180. 10.1038/s41477-018-0116-y

