## Supplemental Information for "Salicylic acid-triggered apoplastic proteolysis releases cryptic phytocytokines with distinct immunogenic functions"

**Supplementary Figures:**
**Figure S1.** Peptide abundance over time after SA treatment.
**Figure S2.** MIF-His protein purification
**Figure S3.** Stability of MIF in different fractions in the presence of PMSF.

**Supplementary Tables:**
**Table S1.** Peptides identified by MS-based peptidomics analysis of maize leaf apoplastic fluid collected 3 h after salicylic acid (SA) or buffer (mock) infiltration
**Table S2.** Peptides identified by MS-based peptidomics analysis of maize leaf apoplastic fluid collected 6 h, 9 h and 12 h after salicylic acid (SA) or buffer (mock) infiltration
**Table S3.** Selected peptides from MS analysis of APF
**Table S4.** Table of differential gene expression analysis.
**Table S5.** Significantly enriched GO terms for Biological Processes (BP) in the upregulated genes at 2 hpi and 24 hpi.
**Table S6.** Peptides identified by MS-based peptidomics analysis of maize leaf apoplastic fluid collected 3h following SA, SA+PMSF or buffer (mock) infiltration
**Table S7.** Cleavage products of ZmMDL1-His (Zm00001eb226780) following co-incubation with fraction 7
**Table S8.** Oligonucleotides

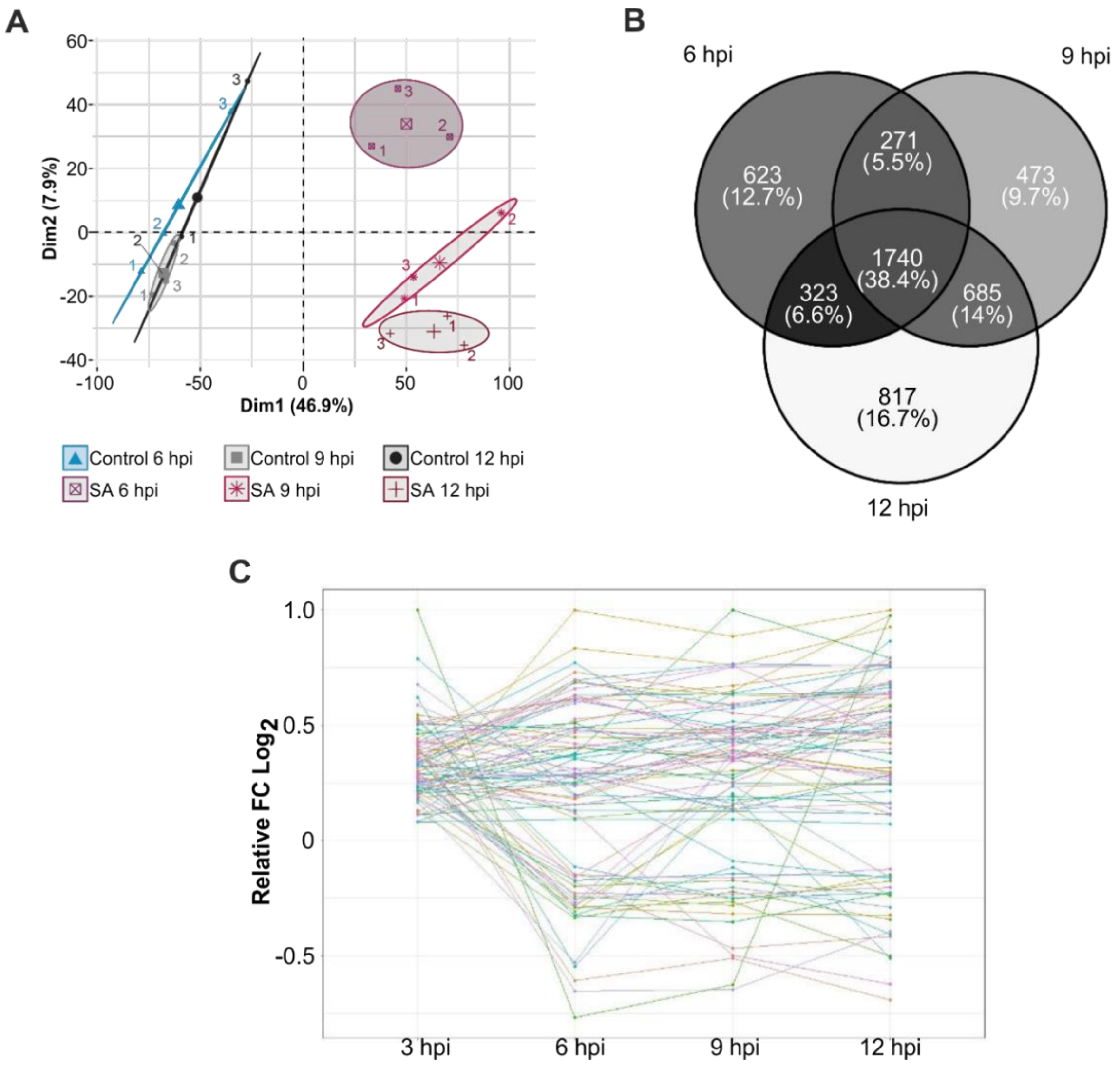


**Fig S1. Peptide abundance over time after SA treatment.** Peptides found via MS analysis in the APF of SA- or control treated maize plants at 6, 9 and 12 hpi. (A) PCA plot of the SA/control depicting the distribution and change of the peptidome over time. (B) Venn diagram of peptides found to be significant after SA treatment at the corresponding time point. For significance students t-test with a permutation-based FDR calculation was performed. The total number of detected peptides for each time point is indicated below each time point (C) Change of abundance of selected peptides over time. Visualized is the relative FC Log2 of the peptides relative to the highest value. Selected were all peptides, which were significantly more abundant at 3 hpi after SA treatment. Significant differences were calculated based on students t-test.

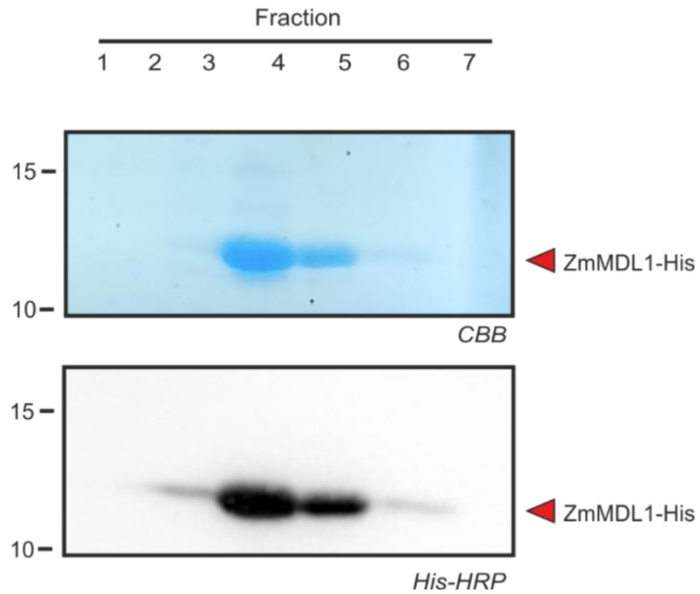


**Fig S2. MIF-His protein purification.** Recombinant ZmMDL1-His was heterologously expressed and purified by immobilized metal affinity chromatography (IMAC) using a 1-mL HisTrap FF Crude column (Cytiva) on an ÄKTA Start chromatography system. Elution fractions were separated by SDS–PAGE and visualized by Coomassie Brilliant Blue (CBB) staining. A prominent band corresponding to ZmMDL1-His was detected in elution fractions 4 and 5 and was subsequently confirmed by anti-His immunoblotting.

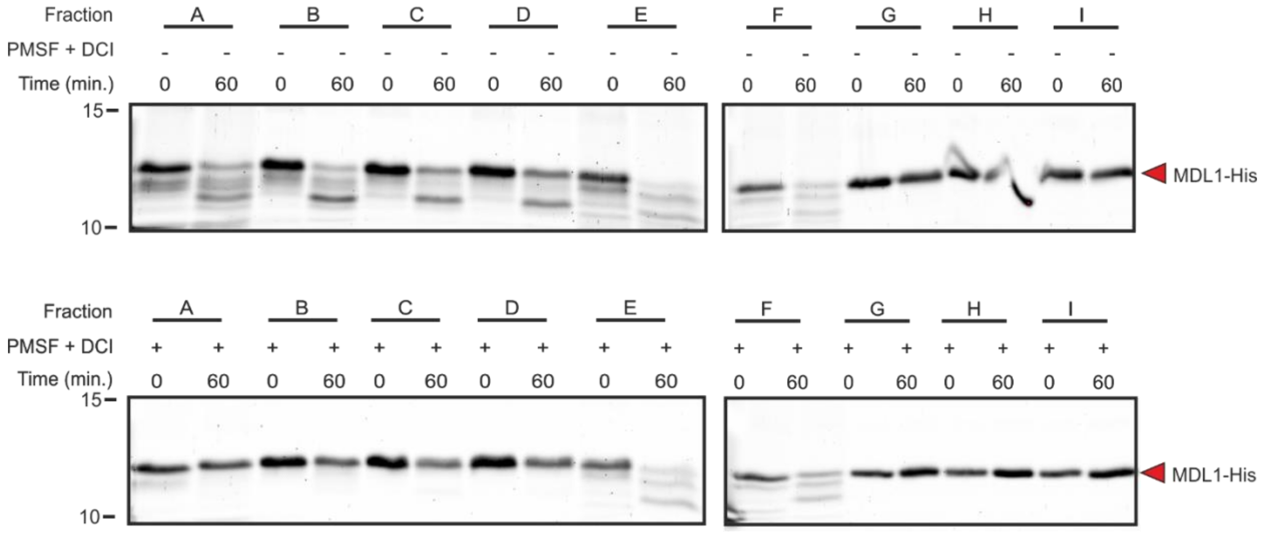


**Fig S3. Stability of ZmMDL1 in different fractions in the presence of PMSF.** Fractions were grouped (A–I), and ZmMDL1 cleavage activity was assessed for each group. Fractions were pre-incubated for 15 min with EDTA (5 mM), pepstatin A (100 µM), and E-64 (100 µM), in the presence or absence of PMSF (100 µM) and DCI (100 µM). Proteins were separated by SDS–PAGE, stained with Sypro Ruby, and analyzed by fluorescence scanning.
